# Cosolvent Effects on the Growth of Protein Aggregates Formed by a Single Domain Globular Protein and an Intrinsically Disordered Protein

**DOI:** 10.1101/490136

**Authors:** Balaka Mondal, Govardhan Reddy

## Abstract

Cosolvents modulate the stability of protein conformations and exhibit contrasting effects on the kinetics of aggregation by globular proteins and intrinsically disordered proteins (IDPs). The growth of ordered protein aggregates, after the initial nucleation step is believed to proceed through a dock-lock mechanism. We have studied the effect of two denaturants (guanidinium chloride (GdmCl) and urea) and four protective osmolytes (trimethylamine N-oxide (TMAO), sucrose, sarcosine, and sorbitol) on the free energy surface (FES) of the dock-lock growth step of protein aggregation using a coarse-grained protein model and metadynamics simulations. We have used the proteins cSrc-SH3 and *Aβ*_9−40_ as model systems representing globular proteins and IDPs, respectively. The effect of cosolvents on protein conformations is taken into account using the molecular transfer model (MTM). The computed FES shows that protective osmolytes stabilize the compact aggregates, while denaturants destabilize them for both cSrc-SH3 and *Aβ*_9−40_. However, protective osmolytes increase the effective energy barrier for the multi-step domain swapped dimerization of cSrc-SH3, which is critical to the growth of protein aggregates by globular proteins, thus slowing down overall aggregation rate. Contrastingly, denaturants decrease the effective barrier height for cSrc-SH3 dimerization, and hence enhances the aggregation rate in globular proteins. The simulations further show that cSrc-SH3 monomers unfold before dimerization and the barrier to monomer unfolding regulates the effective rate of agrgegation. In the case of IDP, *Aβ*_9−40_, protective osmolytes decrease and denaturants increase the effective barriers in the dock-lock mechanism of fibril growth, leading to faster and slower growth kinetics, respectively.

## Introduction

Under physiological conditions, proteins generally exist in native folded form, which are functionally active and carry out a myriad of biological tasks. Upon changes in the physicochemical properties that influence protein stability and dynamics, proteins can undergo misfolding and form aggregates called amyloid fibrils.^1–4^ Misfolded globular proteins and intrinsically disordered proteins (IDPs) form disordered aggregates when their concentration exceeds a critical value. From the disordered protein aggregates, a structured aggregate nucleates and grows through a dock-lock mechanism^5–10^ leading to mature amyloid fibrils, which are associated with a series of neurodegenerative disorders.^11,12^ Initially in the dock stage, the protein in solution interacts with the defects present at the edges of the ordered nucleated fibril and docks onto the fibril. In the lock stage, the protein, which is docked on the fibril, undergoes a structural transition and becomes part of the fibril contributing to its growth.^8–10,13–17^ The structure and growth of amyloid fibrils are affected by several physico-chemical factors like temperature, pressure, pH and cosolvents. An important class of cosolvents are small organic molecules called osmolytes that shifts the equilibrium between folded and unfolded states of the protein,^18–21^ and hence osmolytes can also strongly influence protein aggregation. Numerous experiments^22–33^ reveal contrasting qualitative effects of denaturants and protective osmolytes on the nucleation and growth rate of fibrils by globular proteins and IDPs. A few simulation studies elucidated the effect of cosolvents on the growth of fibrils by small IDPs,^34–36^ and a globular protein.^37^ To understand the mechanistic differences in the fibril growth by globular proteins and IDPs, which will provide insight into the experimental results, we computed the effect of six different cosolvents on the free energy surfaces (FES) for fibril growth using coarse-grained protein models and computer simulations.

Experiments investigating the role of protective osmolytes in aggregation, conclude that generally protective osmolytes delay the aggregation rate in globular proteins,^22–28^ and enhance the aggregation rate in the case of IDPs.^23,29–33^ For a globular protein to aggregate, it has to break a significant fraction of its native contacts present in its folded state, to interact and form inter contacts with another protein present in the solution.^38,39^ The unfolding required for a globular protein to aggregate should favor denaturants in small concentrations over protective osmolytes. Whereas IDPs, which are unstructured in solution, form compact structured aggregates, and this process should favor protective osmolytes over denaturants. It is also difficult to generalize these results for all the proteins, as among other factors, these processes also depend on the sequence details of a protein chain, topology of the folded state and subtle interactions between the protein, solvent and osmolyte.

The effect of cosolvents on the growth rate and aggregate structure depends on the mechanism of aggregation. In the aggregation of globular proteins, experiments^40–46^ and simulations^37^ provide strong evidence for domain swapping, where two or more identical protein chains exchange a part of their secondary structure or a tertiary globular domain. Experiments^47,48^ on human cystatin C show that amyloid formation is suppressed when dimer formation is prevented indicating an interdependence between the two. To explain the role of domain swapping in protein aggregation, a few models are proposed.^43,49^ In the “run away domain swapping” model,^43^ proteins exchange their domains with the neighboring proteins and the chain continues to grow. In the “off-pathway folding” (OFF) model, ^49^ the domain swapped oligomers are proposed to be the end products, which cannot convert to fibrillar structure and they slow down fibril formation. This indicates that domain swapping plays a key role in the aggregation of globular proteins and it is analogous the dock-lock step for fibril growth after the nucleation event.

Experiments and simulations show that domain swapping in globular proteins is governed by the topology of the folded monomers,^53,54^ which undergo at least partial unfolding to initiate domain swapping.^53,55,56^ Studies have shown that a protein monomer can be engineered to undergo domain swapping by modulating the stiffness of the hinge region connecting the two domains in the folded monomer, by introducing mutations in the hinge region, ^56–60^ shortening the hinge region by deleting residues, ^61,62^ lengthening the hinge region by introducing small peptide chains^63^ or proteins,^64^ and by stabilizing the domain swapped dimers by the formation of disulfide bonds.^65^ Simulations^66^ also revealed that functionally active partially folded protein conformations can form domain swapped dimers leading to the hypothesis that domain swapping can be a consequence of protein function imposing constraints on the structrure. Since domain swapping involves protein unfolding, cosolvents, which influence the stability of protein conformations, should also strongly influence domain-swapping in proteins, which is a key step in the growth of aggregates by globular proteins.

In this paper, we studied the effect of four protective osmolytes (TMAO, sucrose, sarcosine, and sorbitol) and two denaturants (urea and GdmCl) on the dock-lock growth step of amyloid fibrils, which contributes to fibril elongation after nucleation. Using cSrc-SH3 and *Aβ*_9−40_ as model systems for globular proteins and IDPs, respectively, we studied the effect of cosolvents on the FES of the dock-lock step (Fig. 1). cSrc-SH3 domain is a ^56^ residue long peptide, which is widely studied both experimentally^51,67–72^ and computationally^73–76^ to probe various aspects of protein folding. This peptide is shown to form intertwined dimers by swapping its RT loop (n-Src loop acting as hinge loop) and amyloid fibrils in mildly acidic conditions.^70^ Whereas *Aβ*_9−40_ peptide is an IDP whose aggregates are associated with neurodegenerative diseases. ^11,12^ It is computationally challenging to construct the FES associated with the fibril elongation process using atomistic description of proteins and unbiased simulations. Coarse-grained protein models^77–84^ have played an important role in elucidating various aspects of nucleation and growth of protein aggregates. To bridge the time scale between simulations and experiments, we modeled the proteins using a coarse-grained self-organized polymer-side chain (SOP-SC) description^75,85^ and used metadynamics simulations^86^ to construct the FES associated with dock-lock step of fibril growth. Co-solvent effects are incorporated in the FES using the molecular transfer model (MTM).^87,88^

**Figure 1:**
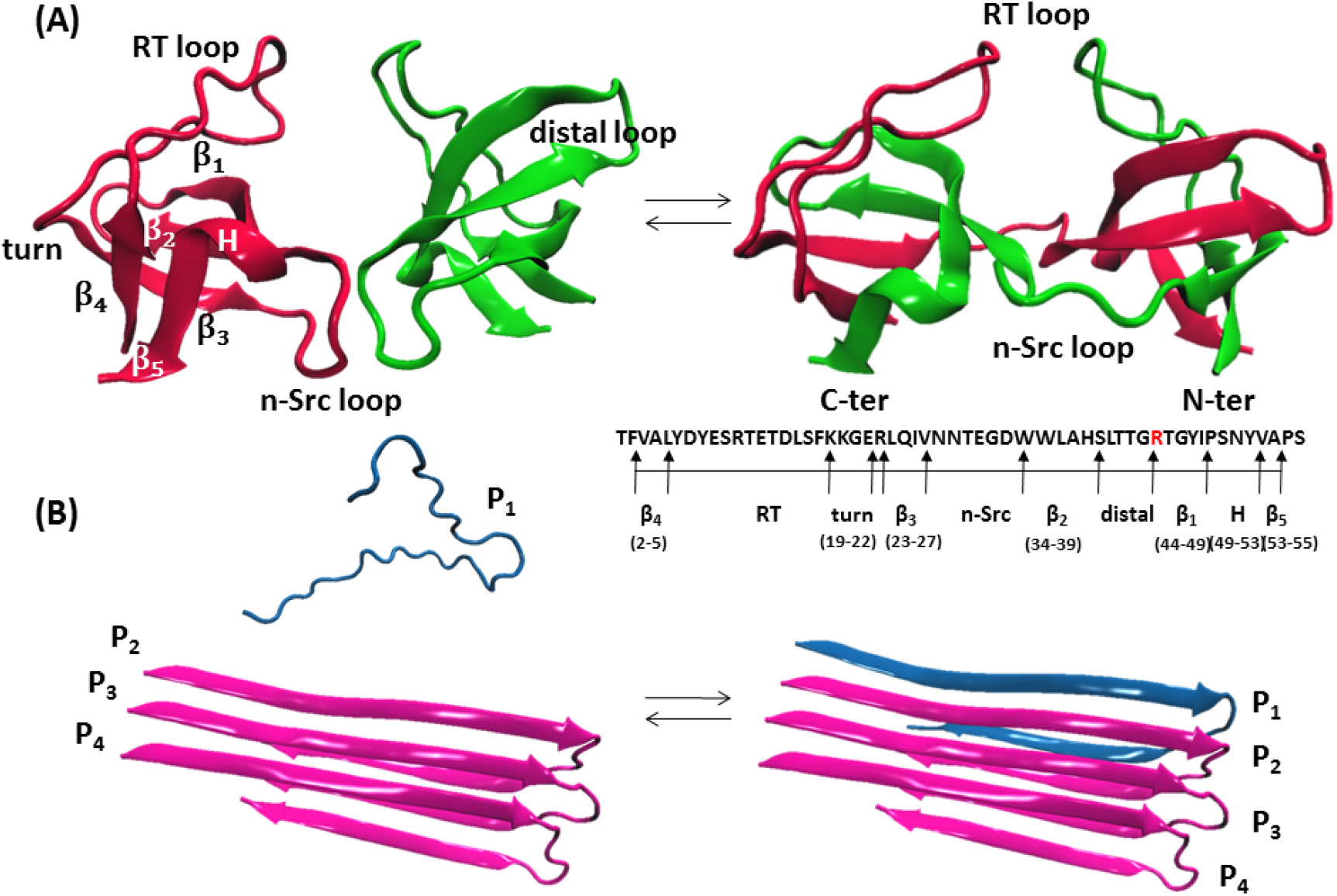
Schematic of the dock-lock step in the growth of fibrils by a globular protein and an IDP. (A) The NMR structure^50^ of SH3 domain of cSrc tyrosine kinase (PDB ID: 1SRL) consists of five beta strands (*β*_1_-*β*_5_), out of which *β*_1_ and *β*_2_ are connected via distal loop, *β*_2_ and *β*_3_ are connected via n-Src loop, and *β*_3_ and *β*_4_ are connected via RT loop and turn. cSrc-SH3 module swaps its RT loop with an identical monomer to form an intertwined dimeric aggregate (PDB ID: 3FJ5).^51^ The monomer has ^56^ residues and the amino acid sequence and corresponding secondary structure is reported below the dimer structure. In the dimer, the glutamine residue in the 44th position is mutated to arginine residue. The free energy is computed for the dimerization of cSrc-SH3 monomers. (B) The amyloid fibril template of the IDP *Aβ*_9−40_ made from three peptides labeled *P*_2_, *P*_3_ and *P*_4_ is constructed using the solid state NMR structure (PDB ID: 2LMN).^52^ The residues 1-8 from the N-terminal in each peptide are disordered in the fibril and are truncated. The free energy is computed for the binding of an unbound *Aβ* peptide (*P*_1_) to the fibril template (*P*_2_ − *P*_4_).

## Results and Discussions

### Free Energy Surface for Fibril Elongation

Experiments^44^ stipulate that domain swapping is a key step in the aggregation of globular proteins. To elucidate the mechanism we computed the underlying FES of domain swapping in a single domain globular protein, cSrc-SH3, which is found to form intertwined domain swapped dimers.^46,51^ We used coarse-grained SOP-SC model of the Q128E mutant of the dimeric crystal structure^51^ (PDB ID: 3FJ5) as a reference system (Fig. 1A). A detailed description of the SOP-SC model and the force-field is described in the Supporting Information (SI). The FES (Fig. 2A) for the domain swapped dimerization of cSrc-SH3 is constructed using a combination of umbrella sampling^89^ and metadynamics86,^90,91^ simulations as described in the SI. The free energy is projected onto three collective variables (CVs): (1) centre of mass distance (*R_cm_*) between the cSrc-SH3 monomers (*M*_1_ and *M*_2_), (2) intra native contacts present in the folded cSrc-SH3 monomers (*Q_intra,dim_*), and (3) inter native contacts present in the domain swapped cSrc-SH3 dimer (*Q_inter_*). For efficient exploration of the underlying energy landscape, we used umbrella sampling on the CV *R_cm_*, where we added a restraint on *R_cm_* using a harmonic potential. To explore all possible scenarios, where the separation between monomers is small enough to facilitate dimerization, to large separation between monomers where dimerization is not possible, we varied *R_cm_* between 2 Å to ^35^ Å using multiple umbrella sampling windows. The difference in *R_cm_* between successive umbrella windows is 1 Å. In each of these umbrella sampling windows, metadynamics simulations are run on the CVs, *Q_intra,dim_* and *Q_inter_*, to compute the FES for domain-swapped dimer formation, *F* (*Q_inter_, Q_intra,dim_ | R_cm_*). The convergence of the metadynamics simulations is shown in Fig. S1 in the SI. Finally the free energy *F* (*R_cm_, Q_inter_, Q_intra,dim_*) is obtained by combining *F* (*Q_inter_, Q_intra,dim_ | R_cm_*) obtained from different umbrella windows using the weighted histogram analysis method (WHAM)^91,92^ (details are in the SI). The free energy *F* (*Q_inter_, Q_intra,dim_*) is computed using the equation

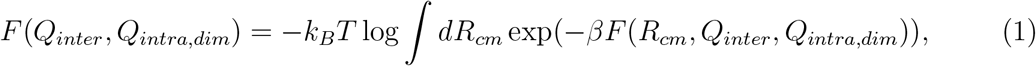

where *β* = 1/*k_B_T*, *k_B_* is the Boltzmann constant and *T* is the temperature. The contour plot of *F* (*Q_inter_, Q_intra,dim_*) for cSrc-SH3 dimer formation is computed at *T* = 353 K, which is 0.96×*T_M_*, where *T_M_* is the melting temperature of the coarse-grained cSrc-SH3 monomer. To facilitate conformational transitions in cSrc-SH3, we chose *T* close to *T_M_*.

The free energy *F* (*Q_inter_, Q_intra,dim_*), shows five major basins denoted as A to E, when the total number of intra native contacts present in both the monomers (*Q_intra,dim_*) are considered (Fig. 2A). The basins correspond to both the monomers folded (basin A), one monomer folded and the second monomer unfolded (basin B), both the monomers unfolded (basin C), intermediate populated during domain swapping (basin D) and domain swapped monomers forming a dimer (basin E). When we project the free energy onto *Q_inter_* and the number of intra native contacts present in only a single monomer (*Q_intra,mon_*), the free energy *F* (*Q_inter_, Q_intra,mon_*) shows only four basins with the basin B missing (Fig. 2B). This proves that indeed, basin B is composed of one folded and one unfolded monomers, and there is no partially folded state involved. The computed FES reveals that unfolded monomers swap their RT loops and undergo partial structuring to give rise to swapped dimers, which is very similar to the domain swapping mechanisms observed in Cyanovirin-N^55^ and p13suc1^56^ proteins. Basin D in both *F* (*Q_inter_, Q_intra,dim_*) and *F* (*Q_inter_, Q_intra,mon_*) appear at similar values of *Q_inter_*, indicating that it is an intermediate in the dimer formation (Fig. 2A, B).

**Figure 2:**
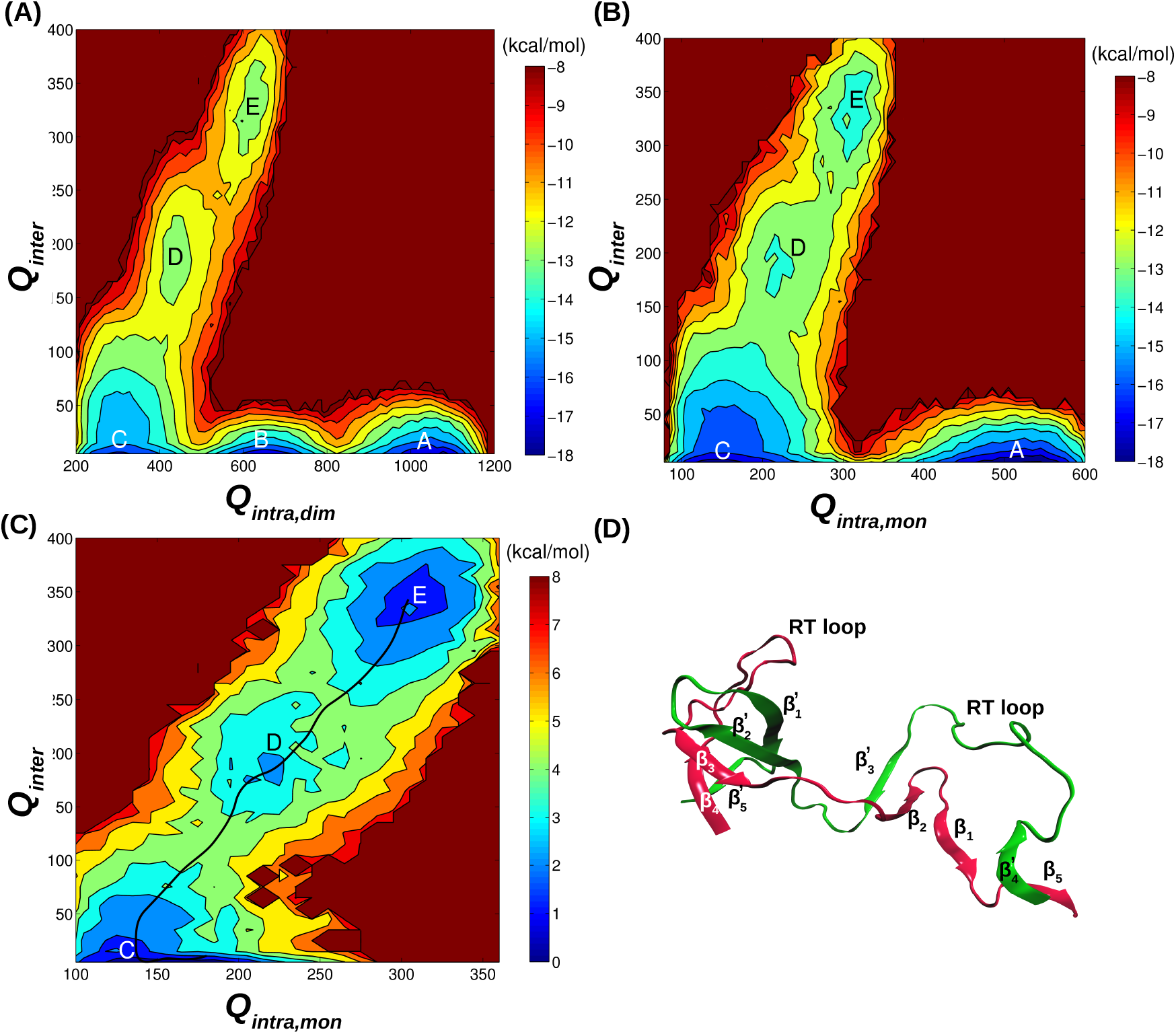
Amyloid fibrils of globular proteins grow by domain-swapping in the dock-lock step. (A) The free energy of cSrc-SH3 domain swapping, *F* (*Q_inter_, Q_intra,dim_*), computed using metadynamics and umbrella sampling simulations, is projected onto the number of inter native contacts present between the two monomers in the domain swapped dimer (*Q_inter_*), and the number of native intra contacts present in both the folded monomers (*Q_intra,dim_*). The free energy landscape shows five basins corresponding to the folded monomers (basin A), an intermediate state where one monomer is folded and the other monomer is unfolded (basin B), both the monomers unfolded (basin C), partially formed dimer (basin D) and domain swapped dimer (basin E). (B) Free energy surface, *F* (*Q_inter_, Q_intra,mon_*), projected onto *Q_inter_* and the number of native intra contacts present in only one monomer (*Q_intra,mon_*). Basin B disappears from the free energy landscape as we eliminate the possibility of one monomer folded and the other monomer unfolded, when we project the free energy onto *Q_inter_* and *Q_intra,mon_*. (C) Free energy surface, *F* (*Q_inter_, Q_intra,mon_ R_cm_* = 4.5 Å), of cSrc-SH3 dimer formation when the center of mass between the two monomer is restrained using a harmonic potential at *R_cm_* = 4.5 Å. The minimum energy path *σ*(*Q_inter_*; *Q_intra,mon_*) corresponding to the formation of the domain swapped dimer (basin E) from the two unfolded protein monomers is shown in black. (D) A representative structure of the partially formed dimer corresponding to basin D. Nearly 33 % of *Q_inter_* contacts are formed in this basin.

To characterize the structure of the dimer intermediate corresponding to basin D, we chose an umbrella window corresponding to *R_cm_* = 4.5 Å, where the distance between the monomers is optimal to form a dimer. The free energy corresponding to this window, *F* (*Q_inter_, Q_intra,mon_ | R_cm_* = 4.5 Å), shows three basins, C, D and E (Fig. 2C). A representative structure of the intermediate from basin D is shown in Fig. 2D. In this intermediate structure, nearly half of the native contacts present in the domain swapped structure are present (Fig. S3). The contact map of the protein conformations in basin D shows that the *β*-strands *β*_2_, *β*_3_ and *β*_4_ in one monomer (*M*_1_) forms contacts with the *β*-strands *β′_3_*, *β′_2_* and *β′_5_*, respectively of the second monomer (*M*_2_) (Fig. 2D and S3). The *RT′* loop of *M*_1_ forms contacts with strand *β′_1_*, of *M*_2_. A closer inspection of the intermediate structure reveals that *RT′* loop of *M*_2_ does not engage in any inter chain interactions in this structure, and this leads to two possibilities. Either the *M*_2_ monomer can swap its *RT′* loop with the *M*_1_ monomer to give rise to a closed dimer (basin E), or it can undergo domain swapping in an end to end fashion with other unfolded monomers as proposed in the “run away domain swapping” model,^43^ resulting in fibrillar aggregates. Thus the intermediate in basin D can act as a template for further aggregation, and in terms of the dock-lock mechanism, basin D and E act as the dock and lock stages, respectively.

The mechanism of domain swapping can also depend on temperature. In this study since we performed simulations close to the melting temperature (*T* = 0.96 *T_M_*), the monomers unfolded completely in the first step of domain swapping as partially folded structures in small globular proteins are not stable at high temperatures. Experiments^93,94^ and simulations^37,95,96^ show that either by introducing mutations or changing the stability conditions by varying temperature and denaturant concentration, or by applying external mechanical force between protein segments, a very low population (< 7%)^93,94^ of transiently folded intermediates can be populated. These transiently populated intermediates, which are not completely unfolded are also shown to form domain swapped structures leading to protein aggregates.^37,94^ As the transiently populated intermediates have very small stabilities it is extremely difficult to compute the free energies for dimerization in these conditions.

We also studied the growth phase in aggregation of intrinsically disordered *Aβ*_9−40_ peptides, where unstructured monomeric *Aβ*_9−40_ peptides in solution, bind to a preformed fibril template yielding *β*-sheet rich aggregates. The SOP-SC structure of the fibril template (Fig. 1B), consisting of three *Aβ*_9−40_ peptide chains (labeled as *P*_2_, *P*_3_ and *P*_4_) is constructed using the solid state NMR structure (PDB ID: 2LMN).^52^ The FES for the binding of an unstructured peptide (*P*_1_) in solution, to the fibril template (*P*_2_ *− P*_4_) is computed using metadynamics simulations (Fig. 1B). During simulations, the fibril template (*P*_2_ − *P*_4_) is kept fixed by applying harmonic restraints on the positions of the coarse-grained beads. The FES is projected onto the CVs, centre of mass distance between the peptides *P*_1_ and *P*_2_ 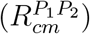, and total number of native contacts, *Q_tot_* (= *Q*_*P*_1__ + *Q*_*P*_2__ + *Q*_*P*_1_, *P*_2__), present when the peptide *P*_1_ is part of the fibril (PDB ID: 2LMN). Here *Q_P_*1 and *Q_P_*2 are the number of intra native contacts present in peptides *P*_1_ and *P*_2_, respectively, and *Q*_*P*_1_, *P*_2__ is the number of inter native contacts present between peptides *P*_1_ and *P*_2_. The convergence of the metadynamics simulations is shown in Fig. S2.

The computed FES *F*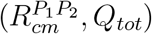 shows two major basins (Fig. 3A). The basin F corresponds to the free unbound state of the peptide *P*_1_, and is characterized by relatively high 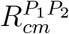 and low values of *Q_tot_*. The basin L corresponds to the locked state where the free monomer *P*_1_ binds to the fibril template and undergoes a structural transition to *β*-strands. In basin L, the peptide *P*_1_ is characterized by low 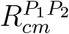 and high *Q_tot_* as it is locked onto the fibril, maximizing the number of contacts with the fibril. Other than these two major basins, a third wide basin is also observed in the vicinity of basin L. This basin corresponds to the docked state (D) of peptide *P*_1_, where it is disordered and partially bound (docked) to the fibril template forming contacts with the peptides from the fibril template. This wide basin indicates that a large number of conformations are possible for the partial binding of the monomer *P*_1_ to the fibril template. This FES strongly supports the dock-lock mechanism of fibril growth.^5–10,14^

**Figure 3:**
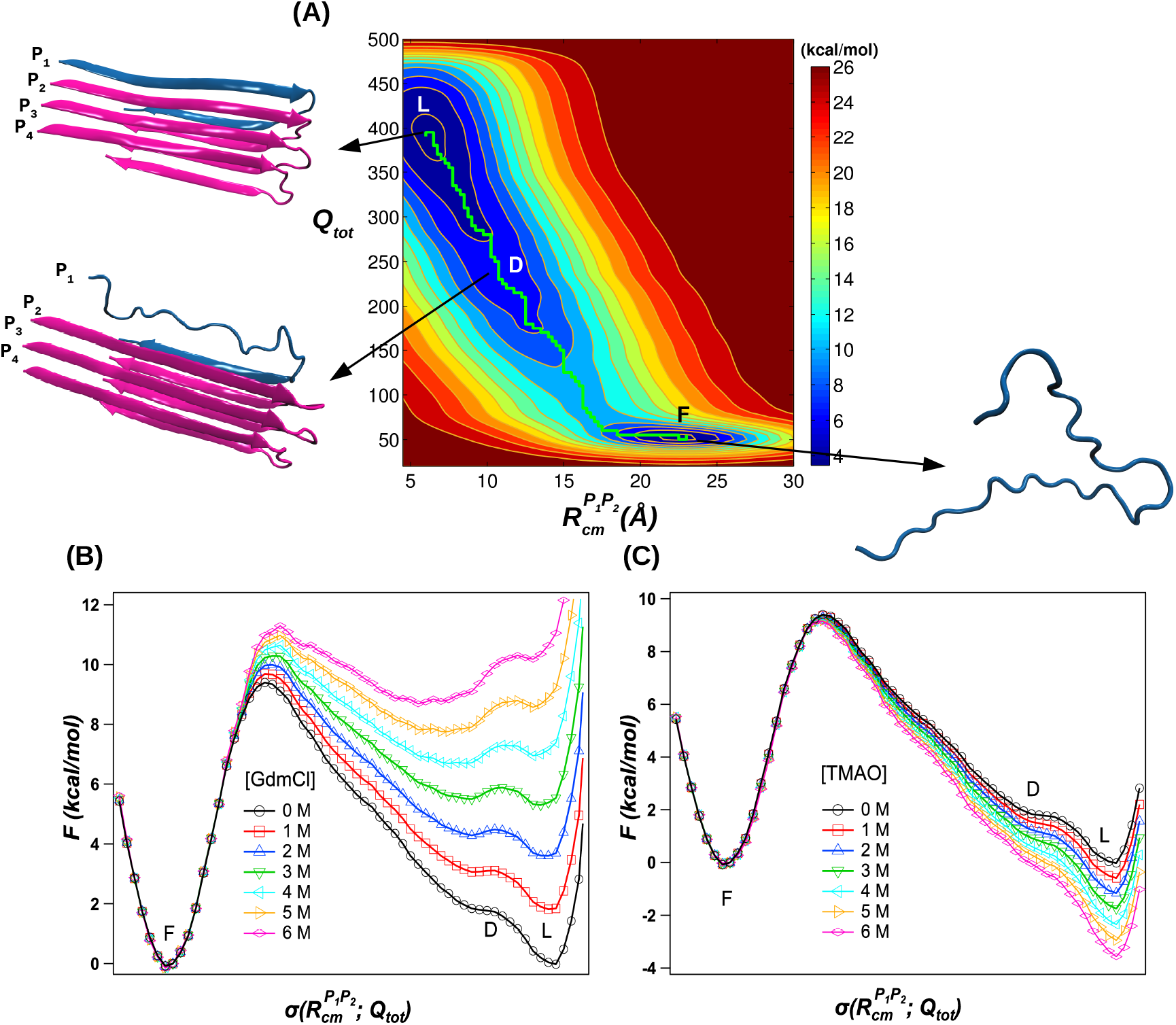
Free energy surface associated with the dock-lock fibril growth step of the IDP *Aβ*_9−40_. (A) The free energy, 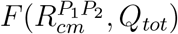, for the binding of a peptide in solution shown in blue (*P*_1_) onto the amyloid fibril template shown in magenta (*P*_2_ - *P*_4_) is projected onto the center of mass distance between the peptides *P*_1_ and 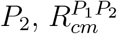, and the total number of native contacts, *Q_tot_*(= *Q*_*P*_1__ + *Q*_*P*_2__ + *Q*_*P*_1_, *P*_2__), where *Q*_*P*_1__, *Q*_*P*_2__ and *Q*_*P*_1_, *P*_2__ are the number of native contacts present in peptides *P*_1_, *P*_2_, and between *P*_1_ and *P*_2_ in the fibril state. The three basins in the free energy correspond to the free state of the monomer *P*_1_ (basin F), the docked state where peptide *P*_1_ interacts with the fibril template (basin D), and the locked state where peptide *P*_1_ becomes part of the fibril (basin L). The free energy is computed at *T* = 350 K and in the absence of cosolvents ([*C*] = 0 M). The minimum energy pathway (MEP), 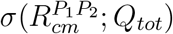, connecting the three basins is shown in green. (B) The effect of denaturant GdmCl on the free energy surface for *Aβ*_9−40_ fibril growth is studied by projecting the free energy onto 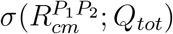. GdmCl stabilizes the free state compared to the docked and locked states, and also increases the barrier height separating the free state from the docked state. (C) The effect of protective osmolyte TMAO on the free energy surface for *Aβ*_9−40_ fibril growth is studied by projecting the free energy onto 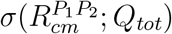. TMAO stabilizes the locked state compared to the free state, and also slightly decreases the barrier height separating the free state and docked state.

### Cosolvent Effects on the Minimum Energy Pathways in the Free Energy Surface

To illustrate the effect of cosolvents on the FES associated with the growth of fibrils formed by both globular proteins and IDPs, we constructed the minimum energy pathway (MEP), connecting various basins on the energy surface^97^ (Fig. 2 and 3). We studied the effect of four different protective osmolytes (TMAO, sucrose, sarcosine and sorbitol), and two different denaturants (urea and GdmCl), on the MEPs. The effect of cosolvents on protein conformations is taken into account using MTM and perturbation theory.^74,87^ The probability of observing protein conformations generated using metadynamics simulations^86,91^ in the absence of a cosolvent ([*C*] = 0) are reweighed^90^ to construct the FES for fibril growth in the presence of a cosolvent with concentration [*C*] (see SI for a detailed description of the simulation method and data analysis).

The 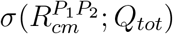 depicting *Aβ*_9−40_ fibril growth is a function of 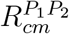 and *Q_tot_*, and connects all the three major basins corresponding to the peptide free state (F), docked state (D) and locked state (L) (Fig. 3A). We probed the effect of cosolvents on fibril growth by studying the variations in the 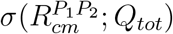 as a function of cosolvent concentration (Fig. 3B, C). In the case of denaturant GdmCl, with increase in GdmCl concentration ([*GdmCl*]), the docked and locked states become unstable compared to the free state, as the denaturants stabilize extended or unfolded states of proteins (Fig. 3B). The barrier height 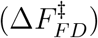 for transition between the free state to the docked state increased by ≈ 2.0 kcal/mol as [*GdmCl*] increased from 0 to 6 M (Table 1). Similar results are also obtained in the case of the denaturant, urea (Fig. S4A). From these results we can conclude that denaturants reduce the growth rate of fibrils formed by IDPs and also destabilize the final aggregate state. In contrast to the denaturants, the protective osmolyte, TMAO, enhanced the stability of the docked and locked state compared to the free state (Fig. 3C and Table 2). There is also a small decrease (≈ 0.3 kcal/mol) in the barrier height 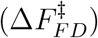 for the transition between the free and docked states indicating that TMAO slightly enhances the growth rates for *Aβ*_9−40_ fibril formation (Table 2). Similar trends are observed in the MEPs in the presence of other protective osmolytes: sucrose, sarcosine and sorbitol (Fig. S4B - S4D), indicating that the protective osmolytes can slightly enhance the fibril growth rates and also stabilize the fibrils formed by IDPs.

**Table 1:**
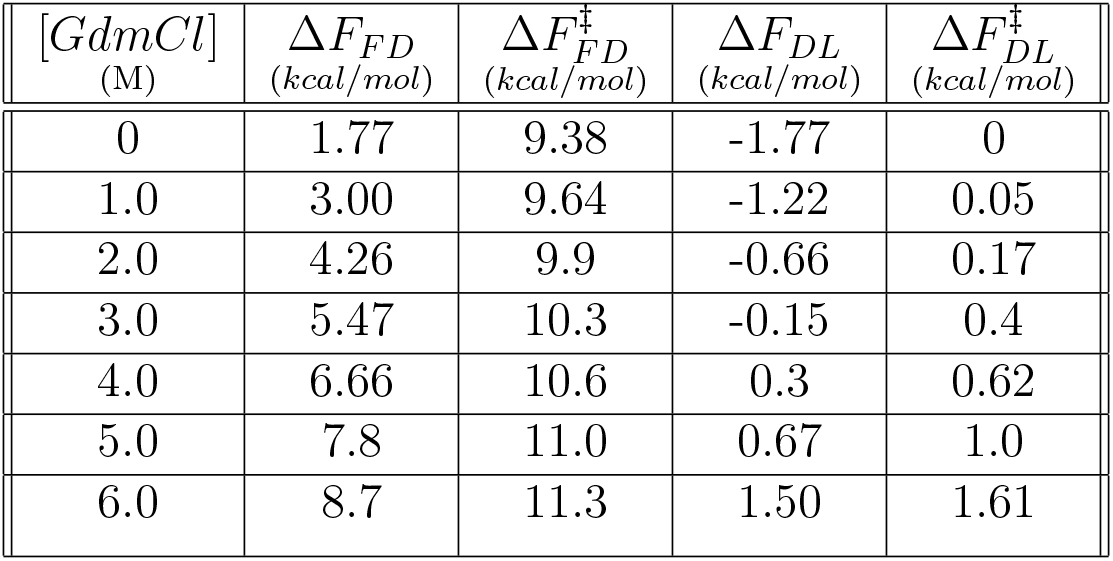
GdmCl effect on the relative difference between the energy minima of different basins, and barrier height separating the basins in the free energy landscape of *Aβ*_9−40_ fibril growth (Fig. 3)

| $[GdmCl]$<br>(M) | $\Delta F_{FD}$<br>(kcal/mol) | $\Delta F_{FD}^{\ddagger}$<br>(kcal/mol) | $\Delta F_{DL}$<br>(kcal/mol) | $\Delta F_{DL}^{\ddagger}$<br>(kcal/mol) |
| --- | --- | --- | --- | --- |
| 0 | 1.77 | 9.38 | -1.77 | 0 |
| 1.0 | 3.00 | 9.64 | -1.22 | 0.05 |
| 2.0 | 4.26 | 9.9 | -0.66 | 0.17 |
| 3.0 | 5.47 | 10.3 | -0.15 | 0.4 |
| 4.0 | 6.66 | 10.6 | 0.3 | 0.62 |
| 5.0 | 7.8 | 11.0 | 0.67 | 1.0 |
| 6.0 | 8.7 | 11.3 | 1.50 | 1.61 |

**Table 2:** TMAO effect on the relative difference between the energy minima of different basins, and barrier height separating the basins in the free energy landscape of *Aβ*_9−40_ fibril growth (Fig. 3)

| $[TMAO]$<br>(M) | $\Delta F_{FD}$<br>(kcal/mol) | $\Delta F_{FD}^{\ddagger}$<br>(kcal/mol) | $\Delta F_{DL}$<br>(kcal/mol) | $\Delta F_{DL}^{\ddagger}$<br>(kcal/mol) |
| --- | --- | --- | --- | --- |
| 0 | 1.77 | 9.38 | -1.77 | 0 |
| 1.0 | 1.47 | 9.34 | -2.10 | - |
| 2.0 | 1.1 | 9.3 | -2.35 | - |
| 3.0 | 0.7 | 9.24 | -2.54 | - |
| 4.0 | 0.3 | 9.20 | -2.77 | - |
| 5.0 | -0.04 | 9.16 | -2.92 | - |
| 6.0 | -0.44 | 9.1 | -3.1 | - |

The effect of cosolvents on domain swapping in globular proteins (cSrc-SH3), which is analogous to the dock-lock step is interesting as domain swapping involves two major steps. In the first step, the proteins unfold increasing the surface area of the protein exposed to solvent. In the second step, unfolded proteins during the course of their folding, domain swap and form dimers, where the total surface area of the proteins exposed to solvent decreases. Denaturants such as urea and GdmCl favor the first step, and hinder the second step in contrast to the protective osmolytes. This implies that both denaturants and protective osmolytes can either increase or decrease the overall transition rate for the protein monomers to form a domain swapped dimer.

The FES for domain swapping in cSrc-SH3 shows that in the first step, the folded protein monomers undergo unfolding, *i.e*. transition from basin A to C (Fig. 2B). The free energy for this transition, when projected onto *Q_intra,mon_*, shows that as expected GdmCl stabilizes the unfolded state compared to the folded state by 2.5 kcal/mol and decreases the barrier height 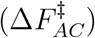 for unfolding by 1.6 kcal/mol on increasing [*GdmCl*] from 0 to 1 M (Fig. 4A and Table 3). This shows that GdmCl stabilizes unfolded state of cSrc-SH3 monomer compared to folded state, and also increases the protein unfolding rate. Whereas on increasing [*TMAO*] from 0 to 6 M, the barrier 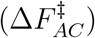 to unfolding increases by 1.3 kcal/mol, and the unfolded state gets destabilized by 2 kcal/mol compared to the folded state (Fig. 4B and Table 4). This shows that TMAO destabilizes the cSrc-SH3 monomer unfolded state compared to folded state, and also decreases the protein unfolding rate.

**Figure 4:**
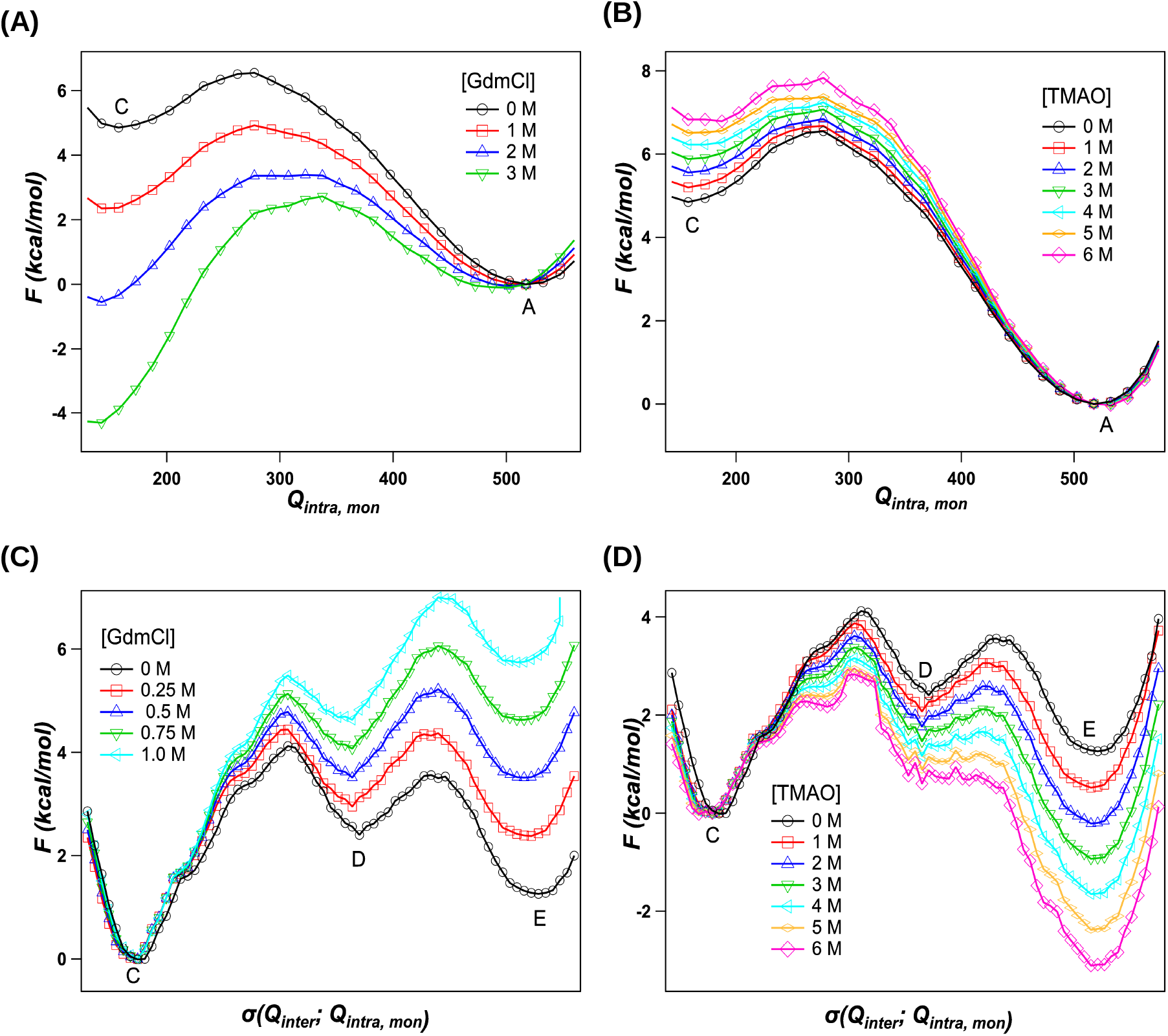
Cosolvent effects on the domain-swapped dimerization of cSrc-SH3. The effect of (A) GdmCl and (B) TMAO on the unfolding of cSrc-SH3 monomer is studied by projecting the free energy *F* (*Q_inter_, Q_intra,mon_ R_cm_* = 32 Å) onto the number of native contacts present in a monomer, *Q_intra,mon_*. GdmCl stabilizes the protein unfolded state compared to the folded state and also decreases the barrier height for transition to the unfolded state from folded state. TMAO destabilizes the protein unfolded state compared to the folded state and increases the barrier height for transition to the unfolded state. The effect of (C) GdmCl and (D) TMAO on the formation of domain swapped dimer from unfolded states is studied by projecting the free energy surface *F* (*Q_inter_, Q_intra,mon_ R_cm_* = 4.5 Å) onto the MEP, *σ*(*Q_inter_*; *Q_intra,mon_*). GdmCl destabilizes the intermediate and dimer corresponding to the basins D and E, respectively, compared to the protein unfolded state in basin C. GdmCl also increases the barrier height for the transition to basins D and E from basin C. Whereas TMAO stabilizes the intermediate and the dimer state compered to the unfolded states and also decreases the height of the barrier for transition to the dimer state from the unfolded state.

**Table 3:**
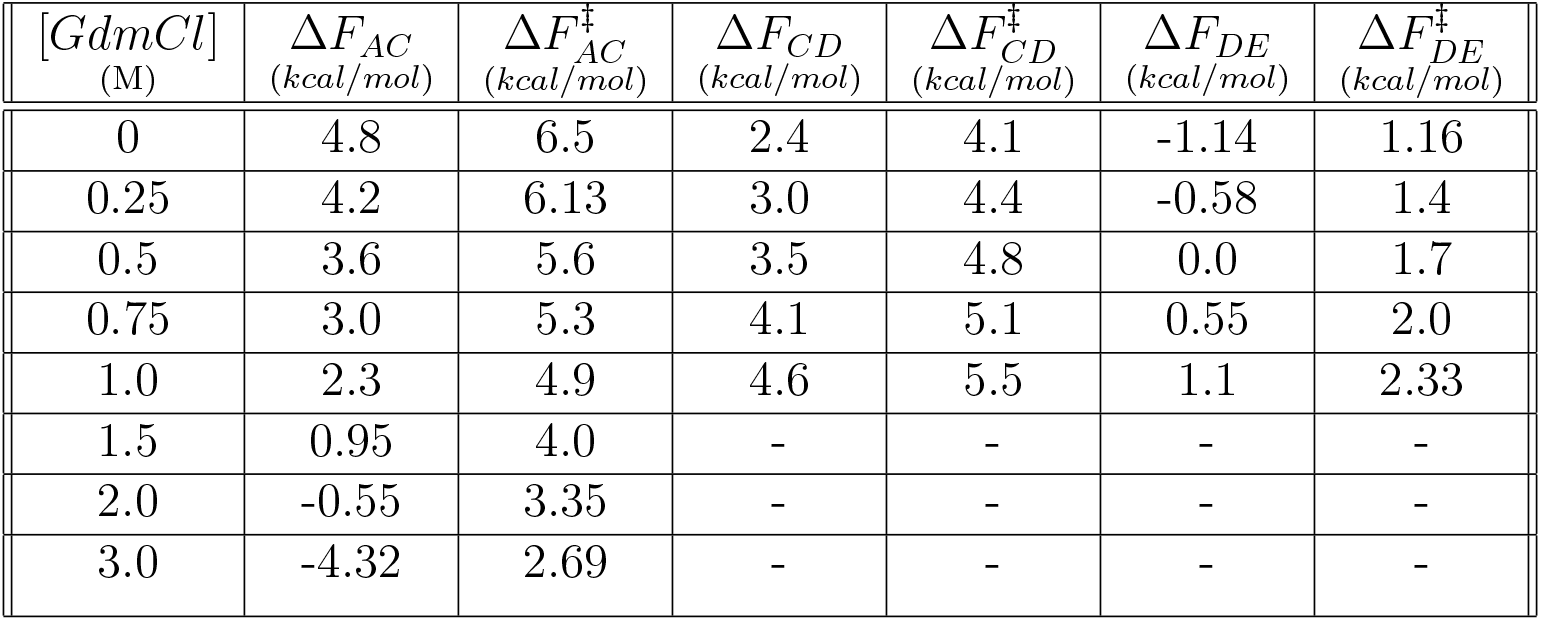
GdmCl effect on the relative difference between the energy minima of different basins, and barrier height separating the basins in the free energy landscape of cSrc-SH3 dimerization (Fig. 2B,C and 4A,C)

To illustrate the effect of cosolvents on the second step in cSrc-SH3 dimer formation, where domain swapping transition takes place from the protein unfolded states (basin C) to the dimer state (basin E), we constructed the MEP *σ*(*Q_inter_*; *Q_intra,mon_*) on the FES *F* (*Q_inter_, Q_intra,mon_ | R_cm_* = 4.5 Å) (Fig. 2C). In contrast to the protein unfolding step, on increasing [*GdmCl*] from 0 to 1 M, the stability of both the intermediate state (basin D) and dimer state (basin E) decreases by 2.2 kcal/mol and 4.4 kcal/mol, respectively, compared to the unfolded state (basin C), while the barrier height for intermediate formation 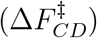, and dimer formation 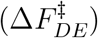, increases by 1.4 kcal/mol and 1.16 kcal/mol, respectively (Fig. 4C and Table 3). This shows that GdmCl not only decreases the stability of the intermediate state and cSrc-SH3 domain swapped dimer compared to the monomer unfolded states, but also it diminishes the growth rate. Similar trends in the FES are also observed for the denaturant, urea (Fig. S5A - S5B). In contrast, on increasing [*TMAO*] from 0 to 6 M, the intermediate state (basin D) and the dimer state (basin E) are stabilized by 1.8 kcal/mol and 4.4 kcal/mol, respectively compared to the unfolded state (basin C), while the barrier height for intermediate formation 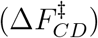, and dimer formation 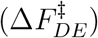 decreases by ≈ 1.3 kcal/mol and 1.0 kcal/mol, respectively (Fig. 4D and Table 4). This implies that TMAO increases the stability of the intermediate state and cSrc-SH3 domain swapped dimer compared to the protein unfolded states, in addition to enhancing the growth rate for intermediate and dimer formation. Similar results are obtained for other protective osmolytes, sucrose, sarcosine and sorbitol (Fig. S5C - S5H).

**Table 4:** TMAO effect on the relative difference between the energy minima of different basins, and barrier height separating the basins in the free energy landscape of cSrc-SH3 dimerization (Fig. 2B,C and 4B,D)

| $[TMAO]$<br>(M) | $\Delta F_{AC}$<br>(kcal/mol) | $\Delta F_{AC}^{\ddagger}$<br>(kcal/mol) | $\Delta F_{CD}$<br>(kcal/mol) | $\Delta F_{CD}^{\ddagger}$<br>(kcal/mol) | $\Delta F_{DE}$<br>(kcal/mol) | $\Delta F_{DE}^{\ddagger}$<br>(kcal/mol) |
| --- | --- | --- | --- | --- | --- | --- |
| 0 | 4.8 | 6.5 | 2.4 | 4.1 | -1.14 | 1.16 |
| 1.0 | 5.17 | 6.7 | 2.0 | 3.8 | -1.55 | 0.98 |
| 2.0 | 5.55 | 6.8 | 1.8 | 3.6 | -2.0 | 0.8 |
| 3.0 | 5.86 | 7.0 | 1.5 | 3.4 | -2.4 | 0.6 |
| 4.0 | 6.2 | 7.2 | 1.2 | 3.1 | -2.8 | 0.45 |
| 5.0 | 6.5 | 7.3 | 0.9 | 2.9 | -3.30 | 0.3 |
| 6.0 | 6.8 | 7.8 | 0.6 | 2.8 | -3.73 | 0.3 |

However, the effect of cosolvents on the rate of domain swapping in globular proteins, which is a key step in the fibril growth is not as trivial as observed in the case of IDPs. The time scale (*τ*) for transition across a barrier of height Δ^*F*‡^ is of the order, *τ* ~ exp(*β*Δ^*F*‡^). Due to the exponential dependence on the barrier height, the effective or overall time scale for dimer formation is strongly influenced by the changes in the largest barrier, which is the transition from basin A to C compared to the transitions between basins C to D, and basins D to E. For example, at *T* = 353 K and [*C*] = 0 M conditions, using the data from Table 4, the ratio of the estimated time scale for transitions between the basins A to C (*τ_AC_*) and basins C to D (*τ_CD_*) is *τ_AC_/τ_CD_* ≈ 30. Since denaturants urea and GdmCl decrease the barrier height between basins A and C, denaturants can effectively increase the rate of cSrc-SH3 domain swapped dimer formation although they destabilize the dimer compared to the folded monomers. On the other hand since protective osmolytes increase the barrier height between basins A and C, they should decrease the effective rate of dimer formation, although they stabilize the dimer state compared to the monomer state. This implies that the growth rate of fibrils formed by globular proteins after nucleation can be enhanced by using denaturants and decreased by protective osmolytes.

The predictions from simulations find support from experimental studies. Experiments studying fibril formation by the globular protein Insulin show that denaturants^24,98^ urea and GdmCl not only decrease the lag time for fibril nucleation, but also increase the fibril elongation rate. Whereas protective osmolytes like TMAO prolonged the nucleation time and also decreased the fibril elongation rate in globular proteins like insulin,^22,23,25,26^ lysozyme,^27^ RNase A^99^ and prion.^28^ In agreement with the predictions from the simulations on IDPs,^34,35^ experiments show that protective osmolytes like TMAO, betaine and sarcosine promote rapid aggregation in the IDPs tau protein,^31,32^ *Aβ*,^29,100^ *α*-synuclein,^23,33^ and hormone glucagon peptide.^30^ Denaturants GdmCl, urea and sodium dodecyl sulfate (SDS) decreased the fibril formation rate in *Aβ* peptides,^36,101^ tau protein^32^ and hIAPP.^102^

In summary, we can conclude that denaturants destabilize and protective osmolytes stabilize the final product, *i.e*. the aggregated state, which is the locked state in the case of the IDP, *Aβ*_9−40_ (Fig. 3), and the domain swapped dimer state in the case of cSrc-SH3 (Fig. 4). The effect of cosolvents on the growth rates of fibrils formed by IDPs is also clear from the free energy surfaces (Fig. 3). GdmCl increased the transition barriers for both the dock and lock steps in the FES, whereas TMAO decreased the barriers. This shows that GdmCl decreases and TMAO increases the fibril growth rate in IDPs. However, the primary finding of this study is the effect of cosolvents on the growth rates of aggregates by globular proteins. Aggregation by globular proteins typically involves domain swapping and using cSrc-SH3 protein as a model system, we showed that this is a mutli-step process. The first step required complete unfolding of cSrc-SH3 monomers, and in the second step these unfolded monomers swapped their *RT* loops and refolded to form domain swapped dimers. Denaturants decreased and increased the barrier height of the first and second steps, respectively; contrarily, osmolytes increased and decreased the barrier height of first and second steps, respectively. However, due to the exponential dependence of rates on barrier heights, the effective rates are mostly governed by the largest barrier and in this case it is the first step, which is the barrier for the unfolding of cSrc-SH3 protein monomers. Thus denaturants enhanced overall rate of dimer formation although they destabilized the dimer compared to the monomers, while protective osmolytes slowed down the growth rate but stabilized the dimers compared to the monomers, and these predictions are in agreement with the experiments. Although we have studied only the effect of cosolvents on the growth step of protein aggregates, these results should also hold for the nucleation step or the lag phase, as unfolding of globular proteins is also a requirement in the nucleation step of protein aggregates when domain swapping is observed.

## Materials and Methods

The coarse-grained SOP-SC models for the dimerization of cSrc-SH3 and fibril growth of *Aβ*_9−40_ are prepared using the structures with PDB ID: 1SRL,^50^ 3FJ5^51^ and 2LMN,^52^ respectively. The detailed description of the coarse-grained model and the energy function is given in the SI. The total energy of a protein conformation in the SOP-SC model described by a set of coordinates, {**r**}, in the absence of a cosolvent ([*C*] = 0) is given by

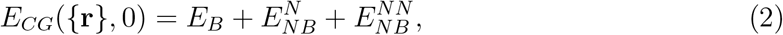

where the term *E_B_* describes the bonded interactions present in the protein, 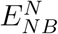 and 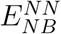 describes the non-bonded native and non-native interactions, respectively. The effect of cosolvent on a protein conformation is taken into account using the molecular transfer model (MTM).^74,87,88^ In the presence of a cosolvent with concentration [*C*], the total energy of a protein conformation^74,87^ with coordinates {**r**} is given by

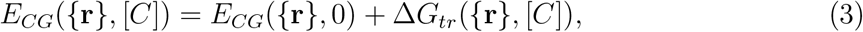

where Δ*G_tr_*({**r**}, [*C*]) is the free energy of transferring a protein conformation with coordinates *{***r***}* from water to a solution with cosolvent concentration [*C*].

Metadynamics simulations^86,90,91^ are performed to compute the free energy for fibril growth in the absence of cosolvents ([*C*] = 0) (see SI for details). To compute the effect of cosolvent on the fibril growth, the term Δ*G_tr_*({**r**}, [*C*]), which describes the effect of co-solvent on a protein conformation is treated as a perturbation in eq. 3. The free energy of fibril growth in the presence of cosolvents is computed by reweighing the probabilities of protein conformations obtained in the absence of cosolvents.^90–92^ Detailed description of the simulation method and data analysis is described in the SI.

## Supporting information

## Acknowledgement

A part of this work is funded by the grants to G.R. from Science and Engineering Research Board (EMR/2016/001356) and Nano mission, Department of Science and Technology, India. (M) acknowledges research fellowship from Indian Institute of Science-Bangalore. The computations are performed using the TUE and Cray XC40 cluster at IISc.

## Supporting Information Available

Detailed description of the protein model, simulation methods, convergence of metadynamics simulations and analysis; Figures S1-S5; This material is available free of charge via the Internet at http://pubs.acs.org/.

## Graphical TOC Entry

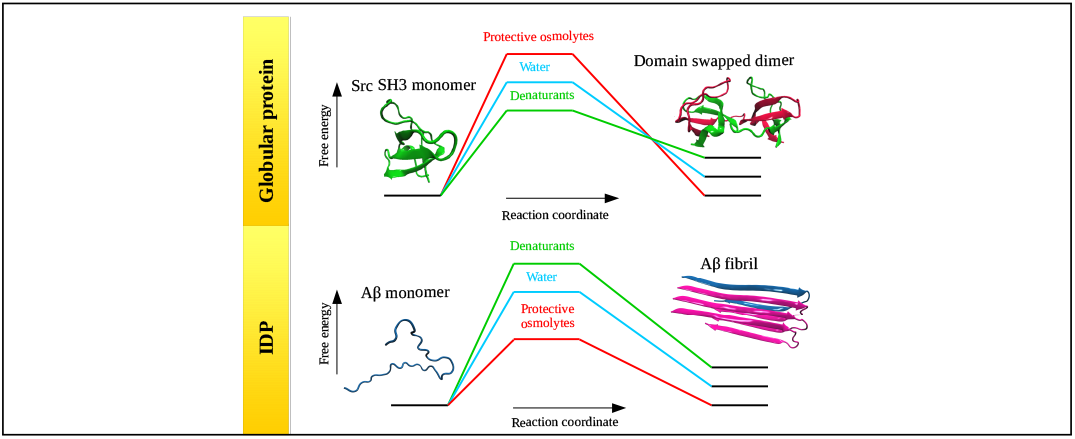

## References

(1) Iadanza, M. G.; Jackson, M. P.; Hewitt, E. W.; Ranson, N. A.; Radford, S. E. A new era for understanding amyloid structures and disease. Nat. Rev. Mol. Cell Biol. 2018, 1–18.

(2) Chiti, F.; Dobson, C. M. Protein Misfolding, Amyloid Formation, and Human Disease: A Summary of Progress Over the Last Decade. Annu. Rev. Biochem. 2017, 86, 27–68.

(3) Straub, J. E.; Thirumalai, D. Toward a molecular theory of early and late events in monomer to amyloid fibril formation. Annu. Rev. Phys. Chem. 2011, 62, 437–463.

(4) Wetzel, R. Kinetics and thermodynamics of amyloid fibril assembly. Acc. Chem. Res. 2006, 39, 671–679.

(5) Esler, W.; Stimson, E.; Jennings, J.; Vinters, H.; Ghilardi, J.; Lee, J.; Mantyh, P.; Maggio, J. Alzheimer’s disease amyloid propagation by a template-dependent dock-lock mechanism. Biochemistry 2000, 39, 6288–6295.

(6) Collins, S. R.; Douglass, A.; Vale, R. D.; Weissman, J. S. Mechanism of Prion propagation: amyloid growth occurs by monomer addition. PLoS. Biol. 2004, 2, 1582–1590.

(7) Cannon, M. J.; Williams, A. D.; Wetzel, R.; Myszka, D. G. Kinetic analysis of beta-amyloid fibril elongation. Anal. Biochem. 2004, 328, 67–75.

(8) Nguyen, P. H.; Li, M. S.; Stock, G.; Straub, J. E.; Thirumalai, D. Monomer adds to preformed structured oligomers of A beta-peptides by a two-stage dock-lock mechanism. Proc. Natl. Acad. Sci. U. S. A. 2007, 104, 111–116.

(9) Reddy, G.; Straub, J. E.; Thirumalai, D. Dynamics of locking of peptides onto growing amyloid fibrils. Proc. Natl. Acad. Sci. U. S. A. 2009, 106, 11948–11953.

(10) O’Brien, E. P.; Okamoto, Y.; Straub, J. E.; Brooks, B. R.; Thirumalai, D. Thermodynamic Perspective on the Dock-Lock Growth Mechanism of Amyloid Fibrils. J. Phys. Chem. B 2009, 113, 14421–14430.

(11) Hardy, J.; Selkoe, D. Medicine - The amyloid hypothesis of Alzheimer’s disease: progress and problems on the road to therapeutics. Science 2002, 297, 353–356.

(12) Lansbury, P. T.; Lashuel, H. A. A century-old debate on protein aggregation and neurodegeneration enters the clinic. Nature 2006, 443, 774–779.

(13) Takeda, T.; Klimov, D. K. Replica exchange simulations of the thermodynamics of a beta fibril growth. Biophys. J. 2009, 96, 442–452.

(14) Rojas, A.; Liwo, A.; Browne, D.; Scheraga, H. A. Mechanism of fiber assembly: treatment of a beta peptide aggregation with a coarse-grained united-residue force field. J. Mol. Biol. 2010, 404, 537–552.

(15) Fawzi, N. L.; Okabe, Y.; Yap, E.-H.; Head-Gordon, T. Determining the critical nucleus and mechanism of fibril elongation of the Alzheimer’s a beta(1–40) peptide. J. Mol. Biol. 2007, 365, 535–550.

(16) Schor, M.; Mey, A. S. J. S.; Noe, F.; MacPhee, C. E. Shedding light on the dock-lock mechanism in amyloid fibril growth using markov state models. J. Phys. Chem. Lett. 2015, 6, 1076–1081.

(17) Han, W.; Schulten, K. Fibril Elongation by Aβ(17–42): Kinetic Network Analysis of Hybrid-Resolution Molecular Dynamics Simulations. J. Am. Chem. Soc. 2014, 136,12450–12460.

(18) Arakawa, T.; Timasheff, S. N. The stabilization of proteins by osmolytes. Biophys. J. 1985, 47, 411–414.

(19) Bolen, D. W. Effects of naturally occurring osmolytes on protein stability and solubility: issues important in protein crystallization. Methods 2004, 34, 312–322.

(20) Canchi, D. R.; Garcia, A. E. Cosolvent effects on protein stability. Annu. Rev. Phys. Chem. 2013, 64, 273–293.

(21) Rani, A.; Venkatesu, P. Changing relations between proteins and osmolytes: a choice of nature. Phys. Chem. Chem. Phys. 2018, 20, 20315–20333.

(22) Choudhary, S.; Kishore, N.; Hosur, R. V. Inhibition of Insulin fibrillation by osmolytes: mechanistic insights. Sci. Rep. 2015, 5, 17599.

(23) White, D. A.; Buell, A. K.; Knowles, T. P. J.; Welland, M. E.; Dobson, C. M. Protein aggregation in crowded environments. J. Am. Chem. Soc. 2010, 132, 5170–5175.

(24) Knowles, T. P. J.; Shu, W.; Devlin, G. L.; Meehan, S.; Auer, S.; Dobson, C. M.; Welland, M. E. Kinetics and thermodynamics of amyloid formation from direct measurements of fluctuations in fibril mass. Proc. Natl. Acad. Sci. U. S. A. 2007, 104,10016–10021.

(25) Arora, A.; Ha, C.; Park, C. Inhibition of Insulin amyloid formation by small stress molecules. FEBS Lett. 2004, 564, 121–125.

(26) Nielsen, L.; Khurana, R.; Coats, A.; Frokjaer, S.; Brange, J.; Vyas, S.; Uversky, V.; Fink, A. Effect of environmental factors on the kinetics of insulin fibril formation: elucidation of the molecular mechanism. Biochemistry 2001, 40, 6036–6046.

(27) Ueda, T.; Nagata, M.; Imoto, T. Aggregation and chemical reaction in hen lysozyme caused by heating at pH 6 are depressed by osmolytes, sucrose and trehalose. J. Biochem. 2001, 130, 491–496.

(28) Tatzelt, J.; Prusiner, S. B.; Welch, W. J. Chemical chaperones interfere with the formation of scrapie Prion protein. Embo J. 1996, 15, 6363–6373.

(29) Yang, D.; Yip, C.; Huang, T.; Chakrabartty, A.; Fraser, P. Manipulating the amyloidbetaaggregationpathway with chemical chaperones. J. Biol. Chem. 1999, 274, 32970–32974.

(30) Macchi, F.; Eisenkolb, M.; Kiefer, H.; Otzen, D. E. The effect of osmolytes on protein fibrillation. Int. J. Mol. Sci. 2012, 13, 3801–3819.

(31) Scaramozzino, F.; Peterson, D. W.; Farmer, P.; Gerig, J. T.; Graves, D. J.; Lew, J. TMAO promotes fibrillization and microtubule assembly activity in the c-terminal repeat region of tau. Biochemistry 2006, 45, 3684–3691.

(32) Levine, Z. A.; Larini, L.; LaPointe, N. E.; Feinstein, S. C.; Shea, J.-E. Regulation and aggregation of intrinsically disordered peptides. Proc. Natl. Acad. Sci. U. S. A. 2015, 112, 2758–2763.

(33) Uversky, V. N.; Li, J.; Fink, A. L. Trimethylamine-N-oxide-induced folding of alpha-synuclein. FEBS Lett. 2001, 509, 31–35.

(34) Muttathukattil, A. N.; Reddy, G. Osmolyte effects on the growth of amyloid fibrils. J. Phys. Chem. B 2016, 120, 10979–10989.

(35) O’Brien, E. P.; Okamoto, Y.; Straub, J. E.; Brooks, B. R.; Thirumalai, D. Thermodynamic perspective on the dock-lock growth mechanism of amyloid fibrils. J. Phys. Chem. B 2009, 113, 14421–14430.

(36) Klimov, D. K.; Straub, J. E.; Thirumalai, D. Aqueous urea solution destabilizes a beta(16–22) oligomers. Proc. Natl. Acad. Sci. U. S. A. 2004, 101, 14760–14765.

(37) Zhuravlev, P. I.; Reddy, G.; Straub, J. E.; Thirumalai, D. Propensity to form amyloid fibrils is encoded as excitations in the free energy landscape of monomeric proteins. J. Mol. Biol. 2014, 426, 2653–2666.

(38) Uversky, V.; Fink, A. Conformational constraints for amyloid fibrillation: the importance of being unfolded. BBA-Proteins Proteomics 2004, 1698, 131–153.

(39) Dobson, C. Protein misfolding, evolution and disease. Trends Biochem. Sci. 1999, 24, 329–332.

(40) Huntington, J.; Pannu, N.; Hazes, B.; Read, R.; Lomas, D.; Carrell, R. A 2.6 angstrom structure of a serpin polymer and implications for conformational disease. J. Mol. Biol. 1999, 293, 449–455.

(41) Janowski, R.; Kozak, M.; Abrahamson, M.; Grubb, A.; Jaskolski, M. 3D domain-swapped human cystatin c with amyloidlike intermolecular beta-sheets. Proteins 2005, 61, 570–578.

(42) Sambashivan, S.; Liu, Y. S.; Sawaya, M. R.; Gingery, M.; Eisenberg, D. Amyloidlike fibrils of ribonuclease a with three-dimensional domain-swapped and native-like structure. Nature 2005, 437, 266–269.

(43) Guo, Z.; Eisenberg, D. Runaway domain swapping in amyloid-like fibrils of T7 endonuclease I. Proc. Natl. Acad. Sci. U. S. A. 2006, 103, 8042–8047.

(44) Bennett, M. J.; Sawaya, M. R.; Eisenberg, D. Deposition diseases and 3D domain swapping. Structure 2006, 14, 811–824.

(45) Liu, C.; Sawaya, M. R.; Eisenberg, D. Beta(2)-microglobulin forms three-dimensional domain-swapped amyloid fibrils with disulfide linkages. Nat. Struct. Mol. Biol. 2011, 18, 49–55.

(46) Bacarizo, J.; Martinez-Rodriguez, S.; Manuel Martin-Garcia, J.; Andujar-Sanchez, M.; Ortiz-Salmeron, E.; Luis Neira, J.; Camara-Artigas, A. Electrostatic effects in the folding of the SH3 domain of the c-Src Tyrosine Kinase: pH-dependence in 3D-domain swapping and amyloid formation. PLoS One 2014, 9.

(47) Nilsson, M.; Wang, X.; Rodziewicz-Motowidlo, S.; Janowski, R.; Lindstrom, V.; Onnerfjord, P.; Westermark, G.; Grzonka, Z.; Jaskolski, M.; Grubb, A. Prevention of domain swapping inhibits dimerization and amyloid fibril formation of cystatin C - Use of engineered disulfide bridges, antibodies, and carboxymethylpapain to stabilize the monomeric form of cystatin C. J. Biol. Chem. 2004, 279, 24236–24245.

(48) Kolodziejczyk, R.; Michalska, K.; Hernandez-Santoyo, A.; Wahlbom, M.; Grubb, A.; Jaskolski, M. Crystal structure of human cystatin C stabilized against amyloid formation. FEBS J. 2010, 277, 1726–1737.

(49) Pallitto, M.; Murphy, R. A mathematical model of the kinetics of beta-amyloid fibril growth from the denatured state. Biophys. J. 2001, 81, 1805–1822.

(50) Yu, H.; Rosen, M. K.; Schreiber, S. L. 1H and 15N assignments and secondary structure of the Src SH3 domain. FEBS Lett. 1993, 324, 87–92.

(51) Camara-Artigas, A.; Martin-Garcia, J. M.; Morel, B.; Ruiz-Sanz, J.; Luque, I. Intertwined dimeric structure for the SH3 Domain of the c-Src tyrosine kinase induced by polyethylene glycol binding. FEBS Lett. 2009, 583, 749–753.

(52) Paravastua, A. K.; Leapman, R. D.; Yau, W. M.; Tycko, R. Molecular structural basis for polymorphism in alzheimer’s beta-amyloid fibrils. Proc. Natl. Acad. Sci. USA 2008, 105, 18349–18354.

(53) Yang, S.; Cho, S.; Levy, Y.; Cheung, M.; Levine, H.; Wolynes, P.; Onuchic, J. Domain swapping is a consequence of minimal frustration. Proc. Natl. Acad. Sci. U. S. A. 2004, 101, 13786–13791.

(54) Mascarenhas, N. M.; Gosavi, S. Understanding protein domain-swapping using structure-based models of protein folding. Prog. Biophys. Mol. Biol. 2017, 128, 113–120.

(55) Liu, L.; Byeon In-Ja, L.; Bahar, I.; Gronenborn, A. M. Domain swapping proceeds via complete unfolding: a F-19- and H-1-NMR study of the Cyanovirin-N protein. J. Am. Chem. Soc. 2012, 134, 4229–4235.

(56) Rousseau, F.; Schymkowitz, J.; Wilkinson, H.; Itzhaki, L. Three-dimensional domain swapping in p13suc1 occurs in the unfolded state and is controlled by conserved proline residues. Proc. Natl. Acad. Sci. U. S. A. 2001, 98, 5596–5601.

(57) Sirota, F. L.; Hêry-Huynh, S.; Maurer-Stroh, S.; Wodak, S. J. Role of the amino acid sequence in domain swapping of the B1 domain of protein G. Proteins 2008, 72,88–104.

(58) Picone, D.; Di Fiore, A.; Ercole, C.; Franzese, M.; Sica, F.; Tomaselli, S.; Mazzarella, L. The role of the hinge loop in domain swapping - The special case of bovine seminal ribonuclease. J. Biol. Chem. 2005, 280, 13771–13778.

(59) Kuhlman, B.; O’Neill, J.; Kim, D.; Zhang, K.; Baker, D. Conversion of monomeric protein L to an obligate dimer by computational protein design. Proc. Natl. Acad. Sci. U. S. A. 2001, 98, 10687–10691.

(60) Murray, A.; Head, J.; Barker, J.; Brady, R. Engineering an intertwined form of CD2 for stability and assembly. Nat. Struct. Biol. 1998, 5, 778–782.

(61) Green, S.; Gittis, A.; Meeker, A.; Lattman, E. One-step evolution of a dimer from a monomeric protein. Nat. Struct. Biol. 1995, 2, 746–751.

(62) Pica, A.; Merlino, A.; Buell, A. K.; Knowles, T. P. J.; Pizzo, E.; D’Alessio, G.; Sica, F.; Mazzarella, L. Three-dimensional domain swapping and supramolecular protein assembly: insights from the X-ray structure of a dimeric swapped variant of human pancreatic RNase. Acta Crystallogr. Sect. D-Struct. Biol. 2013, 69, 2116–2123.

(63) Rowe, C.; Bohm, I.; Thomas, I.; Wilkinson, B.; Rudd, B.; Foster, G.; Blackaby, A.; Sidebottom, P.; Roddis, Y.; Buss, A.; Staunton, J.; Leadlay, P. Engineering a polyketide with a longer chain by insertion of an extra module into the erythromycin-producing polyketide synthase. Chem. Biol. 2001, 8, 475–485.

(64) Ha, J.-H.; Karchin, J. M.; Walker-Kopp, N.; Castaneda, C. A.; Loh, S. N. Engineered Domain Swapping as an On/Off Switch for Protein Function. Chem. Biol. 2015, 22, 1384–1393.

(65) Cho, S.; Levy, Y.; Onuchic, J.; Wolynes, P. Overcoming residual frustration in domain-swapping: the roles of disulfide bonds in dimerization and aggregation. Phys. Biol. 2005, 2, S44–S55.

(66) Mascarenhas, N. M.; Gosavi, S. Protein domain-swapping can be a consequence of functional residues. J. Phys. Chem. B 2016, 120, 6929–6938.

(67) Grantcharova, V. P.; Baker, D. Folding dynamics of the Src SH3 domain. Biochemistry 1997, 36, 15685–15692.

(68) Grantcharova, V.; Riddle, D.; Santiago, J.; Baker, D. Important role of hydrogen bonds in the structurally polarized transition state for folding of the Src SH3 domain. Nat. Struct. Biol. 1998, 5, 714–720.

(69) Grantcharova, V. P.; Baker, D. Circularization changes the folding transition state of the Src SH3 domain. J. Mol. Biol. 2001, 306, 555 – 563.

(70) Bacarizo, J.; Martinez-Rodriguez, S.; Manuel Martin-Garcia, J.; Andujar-Sanchez, M.; Ortiz-Salmeron, E.; Luis Neira, J.; Camara-Artigas, A. Electrostatic effects in the folding of the SH3 domain of the c-Src tyrosine kinase: pH-dependence in 3D-domain swapping and amyloid formation. PLoS One 2014, 9.

(71) Grantcharova, V.; Riddle, D.; Baker, D. Long-range order in the Src SH3 folding transition state. Proc. Natl. Acad. Sci. U.S.A. 2000, 97, 7084–7089.

(72) Martinez, J.; Serrano, L. The folding transition state between SH3 domains is conformationally restricted and evolutionarily conserved. Nat. Struct. Biol. 1999, 6, 1010–1016.

(73) Shea, J. E.; Onuchic, J. N.; Brooks, C. L. Probing the folding free energy landscape of the Src-SH3 protein domain. Proc. Natl. Acad. Sci. U. S. A. 2002, 99, 16064–16068.

(74) Liu, Z.; Reddy, G.; Thirumalai, D. Theory of the molecular transfer model for proteins with applications to the folding of the Src-SH3 domain. J. Phys. Chem. B 2012, 116,6707–6716.

(75) Liu, Z.; Reddy, G.; O’Brien, E. P.; Thirumalai, D. Collapse kinetics and chevron plots from simulations of denaturant-dependent folding of globular proteins. Proc. Natl. Acad. Sci. U. S. A. 2011, 108, 7787–7792.

(76) Ding, F.; Guo, W.; Dokholyan, N. V.; Shakhnovich, E. I.; Shea, J. E. Reconstruction of the Src-SH3 protein domain transition state ensemble using multiscale molecular dynamics simulations. J. Mol. Biol. 2005, 350, 1035 – 1050.

(77) Dima, R.; Thirumalai, D. Exploring protein aggregation and self-propagation using lattice models: Phase diagram and kinetics. Protein Sci. 2002, 11, 1036–1049.

(78) Nguyen, H. D.; Hall, C. K. Molecular dynamics simulations of spontaneous fibril formation by random-coil peptides. Proc. Natl. Acad. Sci. U. S. A. 2004, 101, 16180–16185.

(79) Pellarin, R.; Guarnera, E.; Caflisch, A. Pathways and intermediates of amyloid fibril formation. J. Mol. Biol. 2007, 374, 917–924.

(80) Bellesia, G.; Shea, J.-E. Self-assembly of beta-sheet forming peptides into chiral fibrillar aggregates. J. Chem. Phys. 2007, 126, 245104.

(81) Li, M. S.; Klimov, D. K.; Straub, J. E.; Thirumalai, D. Probing the mechanisms of fibril formation using lattice models. J. Chem. Phys. 2008, 129, 175101.

(82) Auer, S.; Dobson, C. M.; Vendruscolo, M.; Maritan, A. Self-templated nucleation in peptide and protein aggregation. Phys. Rev. Lett. 2008, 101, 258101.

(83) Zhang, J.; Muthukumar, M. Simulations of nucleation and elongation of amyloid fibrils. J. Chem. Phys. 2009, 130, 035102.

(84) Zheng, W.; Tsai, M.-Y.; Chen, M.; Wolynes, P. G. Exploring the aggregation free energy landscape of the amyloid-beta protein (1–40). Proc. Natl. Acad. Sci. U. S. A. 2016, 113, 11835–11840.

(85) Hyeon, C.; Dima, R. I.; Thirumalai, D. Pathways and kinetic barriers in mechanical unfolding and refolding of RNA and proteins. Structure 2006, 14, 1633–1645.

(86) Laio, A.; Parrinello, M. Escaping free-energy minima. Proc. Natl. Acad. Sci. U. S. A. 2002, 99, 12562–12566.

(87) O’Brien, E. P.; Ziv, G.; Haran, G.; Brooks, B. R.; Thirumalai, D. Effects of denaturants and osmolytes on proteins are accurately predicted by the molecular transfer model. Proc. Natl. Acad. Sci. U. S. A. 2008, 105, 13403–13408.

(88) Auton, M.; Bolen, D. W. Predicting the energetics of osmolyte-induced protein folding/unfolding. Proc. Natl. Acad. Sci. U. S. A. 2005, 102, 15065–15068.

(89) Torrie, G.; Valleau, J. Monte-Carlo free-energy estimates using non-Boltzmann sampling - Application to subcritical Lennard-Jones fluid. Chew,. Phys. Lett. 1974, 28,578–581.

(90) Tiwary, P.; Parrinello, M. A time-independent free energy estimator for metadynamics. J. Phys. Chem. B 2015, 119, 736–742.

(91) Awasthi, S.; Kapil, V.; Nair, N. N. Sampling free energy surfaces as slices by combining umbrella sampling and metadynamics. J. Comput. Chem. 2016, 37, 1413–1424.

(92) Kumar, S.; J.M., R.; Bouzida, D.; Swendsen, R.; Kollman, P. The weighted histogam analysis method for free-energy calculations on biomolecules. 1. The method. J. Comput. Chem. 1992, 13, 1011–1021.

(93) Aviram, H. Y.; Pirchi, M.; Barak, Y.; Riven, I.; Haran, G. Two states or not two states: Single-molecule folding studies of protein L. J. Chem. Phys. 2018, 148, 123303.

(94) Neudecker, P.; Robustelli, P.; Cavalli, A.; Walsh, P.; Lundstroem, P.; Zarrine-Afsar, A.; Sharpe, S.; Vendruscolo, M.; Kay, L. E. Structure of an Intermediate State in Protein Folding and Aggregation. Science 2012, 336, 362–366.

(95) Wu, J.; Chen, G.; Zhang, Z.; Zhang, P.; Chen, T. The low populated folding intermediate of a mutant of the Fyn SH3 domain identified by a simple model. Phys. Chem. Chem. Phys. 2017, 19, 22321–22328.

(96) Maity, H.; Reddy, G. Transient intermediates are populated in the folding pathways of single-domain two-state folding protein L. J. Chem. Phys. 2018, 148, 165101.

(97) Marcos-Alcalde, I.; Setoain, J.; Mendieta-Moreno, J. I.; Mendieta, J.; Gomez-Puertas, P. MEPSA: minimum energy pathway analysis for energy landscapes. Bioinformatics 2015, 31, 3853–3855.

(98) Ahmad, A.; Millett, I.; Doniach, S.; Uversky, V.; Fink, A. Partially folded intermediates in insulin fibrillation. Biochemistry 2003, 42, 11404–11416.

(99) Ercole, C.; Pedro Lopez-Alonso, J.; Font, J.; Ribo, M.; Vilanova, M.; Picone, D.; Laurents, D. V. Crowding agents and osmolytes provide insight into the formation and dissociation of RNase A oligomers. Arch. Biochem. Biophys. 2011, 506, 123–129.

(100) Tiiman, A.; Krishtal, J.; Palumaa, P.; Tougu, V. In vitro fibrillization of Alzheimer’s amyloid-beta peptide (1–42). AIP Adv. 2015, 5.

(101) Kim, J.; Muresan, A.; Lee, K.; Murphy, R. Urea modulation of beta-amyloid fibril growth: Experimental studies and kinetic models. Protein Sci. 2004, 13, 2888–2898.

(102) Gao, M.; Winter, R. The effects of lipid membranes, crowding and osmolytes on the aggregation, and fibrillation propensity of human IAPP. J. Diabetes Res. 2015, 2015.

