## Supplementary material for "Cosolvent Effects on the Growth of Protein Aggregates Formed by a Single Domain Globular Protein and an Intrinsically Disordered Protein"

### Methods

#### Self Organized Polymer-Side Chain (SOP-SC) Model for Proteins

We have used the SOP-SC model,[1,2] which is a native-centric coarse-grained protein model in which each amino acid is represented by two beads to model cSrc-SH3 and $A\beta_{9-40}$. The backbone atoms of an amino acid are represented by a bead positioned at the center of $C_\alpha$ atom, and the side chain atoms are represented by a bead positioned at the center of mass of the side-chain. SOP-SC model for monomeric cSrc-SH3, domain swapped cSrc-SH3 and $A\beta_{9-40}$ are constructed using the structures in the protein data bank (PDB) with PDB ID: 1SRL,[3] 3FJ5[4] and 2LMN,[5] respectively. Missing hydrogen atoms are added to the structures using the program visual molecular dynamics (VMD)[6] before calculating the centre of mass of the side chain. The Hamiltonian corresponding to the SOP-SC model is described in terms of bonded ($E_B$) and non-bonded ($E_{NB}$) interactions. Covalently connected beads interact via a bonded potential ($E_B$). Non-bonded interactions ($E_{NB}$) consist of native (N) and non-native (NN) interactions. Interactions between two beads are considered native, if they are separated by at least three bonds and are within a cut-off distance ($R_c$) in the SOP-SC model of the PDB structure. Any other non-covalent interactions are considered as non-native interactions ($E_{NN}$). Native interactions between neighboring side-chain beads are ignored because of their close proximity in both folded and extended unfolded states.

The force-field associated with the SOP-SC model for a protein conformation described by the set of co-ordinates $\{\mathbf{r}\}$ in absence of any cosolvent ($[C] = 0$) is given by

$$E_{CG}(\{\mathbf{r}\}, [0]) = E_B + E_{NB}^{N} + E_{NB}^{NN}. \quad \text{(S1)}$$

The bonds between the beads in the SOP-SC model are modeled using the finite extensible nonlinear elastic (FENE) potential and is given by

$$E_B = -\sum_{i=1}^{N_B} \frac{k}{2} R_0^2 \log\Big(1 - \frac{(r_i - r_{cry,i})^2}{R_0^2}\Big), \tag{S2}$$

where $N_B$ is the total number of bonds between the covalently linked beads in the SOP-SC model, $r_i$ is the distance between $i^{th}$ pair of beads and $r_{cry,i}$ is the distance between the same $i^{th}$ pair of beads in the SOP-SC PDB structure. The values of $k$ and $R_0$ are given in Table S1. The native interactions $E_{NB}^N$, are modeled using a Lennard-Jones type of potential given by

$$\begin{aligned} E_{NB}^N = &\sum_{i=1}^{N_N^{bb}} \epsilon_h^{bb}\big[\big(\frac{r_{cry,i}}{r_i}\big)^{12} - 2\big(\frac{r_{cry,i}}{r_i}\big)^6\big] + \sum_{i=1}^{N_N^{bs}} \epsilon_h^{bs}\big[\big(\frac{r_{cry,i}}{r_i}\big)^{12} - 2\big(\frac{r_{cry,i}}{r_i}\big)^6\big] \\ &+ \sum_{i=1}^{N_N^{ss}} 0.5 \times 300 k_B \times (0.7 - \epsilon_i^{ss})\big[\big(\frac{r_{cry,i}}{r_i}\big)^{12} - 2\big(\frac{r_{cry,i}}{r_i}\big)^6\big], \end{aligned} \tag{S3}$$

where $N_N^{bb}$, $N_N^{bs}$ and $N_N^{ss}$ denote the number of native contact pairs between backbone-backbone, backbone-side chain, and side chain-side chain beads, respectively. $k_B$ is the Boltzmann constant, $r_i$ is the distance between $i^{th}$ pair of beads and $r_{cry,i}$ is the distance between the same $i^{th}$ pair of beads in the SOP-SC PDB structure. The strength of interaction between backbone beads ($\epsilon_h^{bb}$), backbone-side chain beads ($\epsilon_h^{bs}$) are given in Table S1. Values of strength of interaction between side chain beads ($\epsilon_i^{ss}$) are taken from the Betancourt-Thirumalai statistical potential.[7]

Entirely repulsive non-native interactions ($E_{NB}^{NN}$) are modeled as

$$E_{NB}^{NN} = \sum_{i=1}^{N_{NN}} \epsilon_l\big(\frac{\sigma_i}{r_i}\big)^6 + \sum_{i=1}^{N_{ang}^{bb}} \epsilon_l\big(\frac{\sigma^{bb}}{r_i}\big)^6 + \sum_{i=1}^{N_{ang}^{bs}} \epsilon_l\big(\frac{\sigma_i^{bs}}{r_i}\big)^6, \tag{S4}$$

where $N_{NN}$ is the total number of non-native interactions pairs in the SOP-SC model, $\sigma_i$ is sum of the radii of $i^{th}$ pair of beads, and $\sigma^{bb}$ is the diameter of backbone bead. The second and third terms in eq. S4 model the bond angle potential between beads separated by two bonds. $N_{ang}^{bb}$ and $N_{ang}^{bs}$ represent the total number of bond angles between the backbone beads and between the backbone-side chain beads. Values of the side chain radii are given in Table S4. Parameters of the coarse grained energy functions are listed in Table S1.

#### Hamiltonian for the domain swapped dimer of cSrc-SH3

The Hamiltonian for the domain swapped dimerization of two cSrc-SH3 protein monomers (Fig. 1 in the main manuscript) labeled $M_1$ and $M_2$ in absence of any cosolvents ($[C] = 0$) is given by

$$\begin{aligned} E_{M_1,M_2}^{Src}(\{\mathbf{r}\},[0]) = & E_B(M_1) + E_B(M_2) + E_{NB}^{N}(M_1) + E_{NB}^{N}(M_2) \\ & + E_{NB}^{NN}(M_1) + E_{NB}^{NN}(M_2) + E_{NB}^{N}(M_1, M_2) + E_{NB}^{NN}(M_1, M_2), \end{aligned} \tag{S5}$$

where $E_B(M_1)$ and $E_B(M_2)$ represent bonded interactions in the cSrc-SH3 monomers $M_1$ and $M_2$, $E_{NB}^{N}(M_1)$ and $E_{NB}^{N}(M_2)$ represent non-bonded native interactions in $M_1$ and $M_2$, and $E_{NB}^{NN}(M_1)$ and $E_{NB}^{NN}(M_2)$ represent non-bonded non-native interactions in $M_1$ and $M_2$. The native and non-native interactions in cSrc-SH3 protein monomers are extracted using the structure with PDB ID: 1SRL. The terms $E_{NB}^{N}(M_1, M_2)$ and $E_{NB}^{NN}(M_1, M_2)$ in eq. S5 represent the inter native contacts and non-native contacts present between $M_1$ and $M_2$, respectively. These interactions are extracted using the domain swapped dimer structure of cSrc-SH3 with PDB ID: 3FJ5

The inter native interactions between the monomers $M_1$ and $M_2$ is given by

$$\begin{aligned} E_{NB}^{N}(M_1, M_2) = & \sum_{i=1}^{N_N^{bb}(M_1,M_2)} \epsilon_h^{bb}\Big[\Big(\frac{r_{cry,i}}{r_i}\Big)^{12} - 2\Big(\frac{r_{cry,i}}{r_i}\Big)^{6}\Big] + \sum_{i=1}^{N_N^{bs}(M_1,M_2)} \epsilon_h^{bs}\Big[\Big(\frac{r_{cry,i}}{r_i}\Big)^{12} - 2\Big(\frac{r_{cry,i}}{r_i}\Big)^{6}\Big] \\ & + \sum_{i=1}^{N_N^{ss}(M_1,M_2)} 0.5 \times 300 k_B \times (0.7 - \epsilon_i^{ss})\Big[\Big(\frac{r_{cry,i}}{r_i}\Big)^{12} - 2\Big(\frac{r_{cry,i}}{r_i}\Big)^{6}\Big], \end{aligned} \tag{S6}$$

where $N_N^{bb}(M_1, M_2)$, $N_N^{bs}(M_1, M_2)$ and $N_N^{ss}(M_1, M_2)$ represent inter native backbone-backbone, backbone-side chain, and side chain-side chain interactions, respectively. The inter non-native interactions between $M_1$ and $M_2$ are given by

$$E_{NB}^{NN}(M_1, M_2) = \sum_{i=1}^{N_{NN}(M_1,M_2)} \epsilon_l \Big(\frac{\sigma_i}{r_i}\Big)^6, \tag{S7}$$

where $N_{NN}(M_1, M_2)$ is the total number of inter non-native interactions present between the beads of $M_1$ and $M_2$ in the SOP-SC model of the domain swapped structure. The SOP-SC model parameters to study domain swapped dimerization of cSrc-SH3 are given in Table S2.

#### Hamiltonian for the binding of an $A\beta$ peptide in solution to a protofibril

To study the binding of an $A\beta$ peptide in solution to a structured protofibril, we constructed a protofibril made of 3 $A\beta$ peptides using the structure with PDB ID: 2LMN.[5] The protofibril made from the peptides labeled $P_2$, $P_3$, and $P_4$ in Fig. 1 in the main manuscript are restrained to their positions using a harmonic potential. The Hamiltonian for the binding of the peptide $P_1$ in solution to the protofibril in the absence of any cosolvent ($[C] = 0$) is given by

$$\begin{aligned} E_{CG}^{A\beta}(\{\mathbf{r}\}, [0]) = E_B(P_1) + E_B(P_2) + E_B(P_3) + E_B(P_4) + E_{NB}^{N}(P_1) + E_{NB}^{NN}(P_1) \\ + E_{NB}^{N}(P_1, P_2) + E_{NB}^{NN}(P_1; P_2, P_3, P_4) + E_{harm}^{rest}(P_2, P_3, P_4), \end{aligned} \tag{S8}$$

where $E_B(P_1)$, $E_B(P_2)$, $E_B(P_3)$ and $E_B(P_4)$ represent bonded interactions in $P_1$, $P_2$, $P_3$ and $P_4$, respectively, $E_{NB}^{N}(P_1)$ and $E_{NB}^{NN}(P_1)$ represent the native and non-native interactions present within the monomer $P_1$ in the fibril bound state, $E_{NB}^{N}(P_1, P_2)$ are the inter native interactions present between $P_1$ and $P_2$, $E_{NB}^{NN}(P_1; P_2, P_3, P_4)$ represents the repulsive inter non-native interactions present between $P_1$, and the protofibril peptides $P_2$, $P_3$ and $P_4$. The protofibril peptides are restrained to their position using a harmonic potential,

$E^{rest}_{harm}(P_2, P_3, P_4)$. The inter native interactions between $P_1$ and $P_2$ is given by

$$E^{N}_{NB}(P_1, P_2) = \sum_{i=1}^{N^{bb}_N(P_1,P_2)} \epsilon^{bb}_h \big[\big(\frac{r_{cry,i}}{r_i}\big)^{12} - 2\big(\frac{r_{cry,i}}{r_i}\big)^{6}\big] + \sum_{i=1}^{N^{bs}_N(P_1,P_2)} \epsilon^{bs}_h \big[\big(\frac{r_{cry,i}}{r_i}\big)^{12} - 2\big(\frac{r_{cry,i}}{r_i}\big)^{6}\big]$$
$$+ \sum_{i=1}^{N^{ss}_N(P_1,P_2)} 0.5 \times 300 k_B \times (0.7 - \epsilon^{ss}_i)\big[\big(\frac{r_{cry,i}}{r_i}\big)^{12} - 2\big(\frac{r_{cry,i}}{r_i}\big)^{6}\big], \tag{S9}$$

where $N^{bb}_N(P_1, P_2)$, $N^{bs}_N(P_1, P_2)$ and $N^{ss}_N(P_1, P_2)$ are the number of inter native backbone-backbone, backbone-side chain and side chain-side chain interaction pairs, respectively present between $P_1$ and $P_2$. The inter non-native interaction potential between $P_1$ and the peptides in the protofibril is given by

$$E^{NN}_{NB}(P_1; P_2, P_3, P_4) = \sum_{i=1}^{N_{NN}(P_1;P_2,P_3,P_4)} \epsilon_l \big(\frac{\sigma_i}{r_i}\big)^6, \tag{S10}$$

where $N_{NN}(P_1; P_2, P_3, P_4)$ is the total number of inter non-native interactions present between $P_1$ and the protofibril peptides. During the simulation the protofibril is restrained using a harmonic potential given by

$$E^{rest}_{harm}(P_2, P_3, P_4) = \frac{1}{2} k_{harm} \sum_{i=1}^{N_{tot}(P_2,P_3,P_4)} (r_i - r^{ref}_i)^2, \tag{S11}$$

where $N_{tot}(P_2, P_3, P_4)$ is the total number of beads present in the fibril template, $r_i$ and $r^{ref}_i$ are the current and reference position of the $i^{th}$ bead, and $k_{harm}$ is the spring constant. Reference bead positions, $r^{ref}_i$, are obtained from the SOP-SC model of the PDB structure (2LMN). Additionally, we put a harmonic restraint between the center of mass of the peptides $P_1$ and $P_2$ ($R^{P_1P_2}_{cm}$), so that the peptide $P_1$ does not move beyond 50 Å from the protofibril surface and this restraining potential is given by

$$E_{bound} = \begin{cases} \frac{1}{2} k_h (R^{P_1P_2}_{cm} - 50)^2, & R^{P_1P_2}_{cm} > 50 \text{ Å} \\ 0, & R^{P_1P_2}_{cm} \leq 50 \text{ Å}, \end{cases} \tag{S12}$$

where $k_h$ is the spring constant. All the SOP-SC model parameters are given in Table S3.

#### Metadynamics Simulations to Compute Free Energy Surface for Fibril Growth

Computing free energy surfaces for complex multistep processes associated with biomolecules such as aggregation is computationally very demanding using unbiased sampling techniques even with coarse-grained models. We have used an enhanced sampling method called metadynamics[8] and its variants to compute the free energy surface for fibril growth by a globular protein and an IDP.

We have used non-tempered metadynamics simulations to compute the free energy surface of $A\beta$ peptide fibril growth. The free energy is projected onto two collective variables (CVs) by adding the metadynamics bias along these two CVs. The first CV is the center of mass distance between the peptides $P_1$ and $P_2$ ($R_{cm}^{P_1P_2}$), and the second CV is the total contact number of native contacts, $Q_{tot}$ ($= Q_{P_1} + Q_{P_2} + Q_{P_1,P_2}$), present when the peptide $P_1$ is part of the fibril (PDB ID: 2LMN). Here $Q_{P_1}$ and $Q_{P_2}$ are the number of intra native contacts present in peptides $P_1$ and $P_2$, respectively, and $Q_{P_1,P_2}$ is the number of inter native contacts present between peptides $P_1$ and $P_2$. A contact between a pair of beads is computed using the equation

$$Q = \left\{ \frac{1.0 - (\frac{r_i - r_{cry,i}}{r_0})^6}{1.0 - (\frac{r_i - r_{cry,i}}{r_0})^{12}} \right\}, \tag{S13}$$

where $r_i$ is the distance between the $i^{th}$ pair of beads, which are in native contact with each other, $r_0$ is the cutoff distance, and $r_{cry,i}$ is the distance between the same pair of beads in the PDB structure. In metadynamics simulations, in the absence of a cosolvent ($[C] = 0$ M), the system evolves under a time-dependent Hamiltonian given by

$$E_{CG,biased}^{A\beta}\big(\{\mathbf{r}\}, [0], t\big) = E_{CG}^{A\beta}\big(\{\mathbf{r}\}, [0]\big) + V^b\big(R_{cm}^{P_1P_2}, Q_{tot}, t\big), \tag{S14}$$

where $V^b\big(R_{cm}^{P_1P_2}, Q_{tot}, t\big)$ is the metadynamics bias added to the system and it is given by

$$V^b\big(R_{cm}^{P_1P_2}, Q_{tot}, t\big) = \sum_{\tau<t} w \exp\Bigg(-\Bigg[\frac{\big(R_{cm}^{P_1P_2} - R_{cm}^{P_1P_2}(\tau)\big)^2}{2(\delta R_{cm}^{P_1P_2})^2} + \frac{(Q_{tot} - Q_{tot}(\tau))^2}{2(\delta Q_{tot})^2}\Bigg]\Bigg), \quad \text{(S15)}$$

where $\tau$ is the gaussian deposition rate, $\delta R_{cm}^{P_1P_2}$ and $\delta Q_{tot}$ are the gaussian widths and $w$ is the height of the gaussian, which is kept fixed in non-tempered metadynamics. Parameters for metadynamics simulations for $A\beta$ aggregation are listed in Table S5.

To compute the free energy surface for the domain swapped dimerization of cSrc-SH3 we used a technique, which is a combination of non-tempered metadynamics and umbrella sampling simulations developed by Awasthi *et al.*[9] One of the CVs onto which the free energy is projected is the center of mass distance between the cSrc-SH3 monomers $M_1$ and $M_2$ ($R_{cm}$). Using umbrella sampling simulations, the monomers are held at a particular separation, $R_{cm}$, using a harmonic restraint potential given by

$$W_h^b(R_{cm}) = \frac{1}{2}k_h(R_{cm} - R_{cm}^h)^2, \quad \text{(S16)}$$

where $k_h$ is the spring constant, $R_{cm}^h$ is the center of an umbrella window, and the umbrella's were placed along $R_{cm}$ starting from 2.0 Å to 35 Å with a separation of 1.0 Å. In each umbrella window, non-tempered metadynamics simulations were run along two other CVs, $Q_{inter}$ and $Q_{intra,dim}$, to ensure an efficient exploration of the free energy surface. A contact between a pair of bead is computed using eq. S13.

The time dependent Hamiltonian to compute the free energy surface for the domain swapped dimerization of cSrc-SH3 using metadynamics simulations in a particular umbrella sampling window (say $h$) in the absence of any cosolvent ($[C] = 0$ M) is given by

$$E_{CG,M_1,M_2,biased}^{Src}(\{\mathbf{r}\}, [0], t) = E_{CG,M_1,M_2}^{Src}(\{\mathbf{r}\}, [0]) + V_h^b(Q_{inter}, Q_{intra,dim}, t) + W_h^b(R_{cm}),\ h = 1, .....M. \quad \text{(S17)}$$

The metadynamics bias potential $V_h^b(Q_{inter}, Q_{intra,dim}, t)$ is given by

$$V_h^b(Q_{inter}, Q_{intra,dim}, t) = \sum_{\tau<t} w \exp\left( - \left[ \frac{\big(Q_{inter} - Q_{inter}(\tau)\big)^2}{2(\delta Q_{inter})^2} + \frac{\big(Q_{intra,dim} - Q_{intra,dim}(\tau)\big)^2}{2(\delta Q_{intra,dim})^2} \right] \right), \tag{S18}$$

where $\tau$ is the gaussian deposition rate, $\delta Q_{inter}$ and $\delta Q_{intra,dim}$ are the gaussian widths along $Q_{inter}$ and $Q_{intra,dim}$ respectively, $w$ is the constant gaussian height. Parameters for the metadynamics simulations to compute Src SH3 domain swapped dimerization are listed in Table S5.

We performed low-friction Langevin dynamics simulations using the Hamiltonians given by eq. S14 and S17 to compute the free energy surface for fibril growth of cSrc-SH3 and $A\beta_{9-40}$, respectively. The equation of motion in Langevin dynamics is given by

$$m\ddot{\vec{r_i}} = -\zeta\dot{\vec{r_i}} + \vec{F_c} + \vec{\Gamma}, \tag{S19}$$

where $m$ is the mass of a protein bead, $\zeta$ is the friction coefficient, $\vec{r_i}$ is the position of $i^{th}$ bead, $\vec{F_c} = -\frac{\partial E_{CG}^{Met}(\{\mathbf{r}\},0,t)}{\partial \vec{r_i}}$, $\vec{\Gamma}$ is the random force with a white noise spectrum. $E_{CG}^{Met}(\{\mathbf{r}\}, 0, t)$ is the time-dependent Hamiltonian for fibril growth given by eq. S14 and S17. The autocorrelation function of the random force in the discretized form is given by $\langle \Gamma(t)\ \Gamma(t+nh) \rangle = \frac{2\zeta k_B T}{h}\delta_{0,n}$, where $n = 0, 1, ...$ and $\delta_{0,n}$ is the Kronecker delta function. Velocity Verlet algorithm[10,11] is used to integrate the equation of motion. We used $\zeta = 0.05\ m/\tau_L$ and time step, $h = 0.005$ $\tau_L$, where $\tau_L$ is the unit of time used to advance the simulation.

#### Cosolvent Effects and Molecular Transfer Model

We mimicked the effect of cosolvents on protein conformations using the molecular transfer model (MTM).[12–14] In the presence of cosolvents with concentration $[C]$, the time dependent Metadynamics simulation Hamiltonian for the computation of the free energy surface of $A\beta$

peptide binding to a protofibril is given by

$$E^{A\beta}_{CG,biased}\big(\{\mathbf{r}\},[C],t\big) = E^{A\beta}_{CG,biased}\big(\{\mathbf{r}\},[0],t\big) + \Delta G_{tr}(\{\mathbf{r}\},[C]), \tag{S20}$$

and the Hamiltonian for the computation of the free energy surface for domain swapped dimerization of cSrc SH3 is given by

$$E^{Src}_{CG,M_1,M_2,biased}(\{\mathbf{r}\},[C],t) = E^{Src}_{CG,M_1,M_2,biased}(\{\mathbf{r}\},[0],t) + \Delta G_{tr}(\{\mathbf{r}\},[C]), \tag{S21}$$

where $\Delta G_{tr}(\{\mathbf{r}\},[C])$ is the change in free energy on transferring a polypeptide chain with a specific conformation $\{\mathbf{r}\}$ from water to a cosolvent solution with concentration $[C]$, and is given by

$$\begin{aligned}\Delta G_{tr}(\{\mathbf{r}\},[C]) = &\sum_{k=1}^{N_{res}} \delta g^{SC}_{tr,k}([C])\alpha^{SC}_{k}(\{\mathbf{r}\})/\alpha^{SC}_{Gly-k-Gly} \\ &+ \sum_{k=1}^{N_{res}} \delta g^{BB}_{tr}([C])\alpha^{BB}_{k}(\{\mathbf{r}\})/\alpha^{BB}_{Gly-k-Gly},\end{aligned} \tag{S22}$$

where $N_{res}$ represents the total number of residues present in the polypeptide chain, $\delta g^{BB}_{tr}([C])$ and $\delta g^{SC}_{tr,k}([C])$ are the transfer free energies of backbone beads, and $k^{th}$ residue side chain beads, respectively from water to a cosolvent solution with concentration $[C]$, while $\alpha^{BB}_{k}(\{\mathbf{r}\})$ and $\alpha^{SC}_{k}(\{\mathbf{r}\})$ are the solvent accessible surface area (SASA) of the backbone and side chain beads, respectively of the $k^{th}$ residue in the polypeptide chain $\{\mathbf{r}\}$. $\alpha^{BB}_{Gly-k-Gly}$ and $\alpha^{SC}_{Gly-k-Gly}$ represent the SASA values of the backbone and side chain beads of the same amino acid $k$ in the tripeptide $Gly-k-Gly$. The van der Waals radius of the water molecule used in SASA calculation is 1.4 Å. The radii for side chains of amino acids needed to compute $\alpha_k(\{\mathbf{r}\})$ are given in Table S2 in Ref.[15] The transfer free energies,[12,14,16,17] $\delta g_{tr,k}([C])$, for backbone and side chains are listed in Table S3 in Ref.[2] The values for $\alpha_{Gly-k-Gly}$ are listed in Table S4 in Ref.[2]

#### Free Energy Surface for the Binding of an $A\beta$ Peptide in a Cosolvent Solution to a Protofibril

We constructed the time independent free energy landscape of for the binding of $A\beta$ peptide to a protofibril, and cSrc-SH3 domain swapped dimerization using a reweighting technique by Tiwary and Parrinello.[18] The free energy surface for $A\beta$ peptide binding to a protofibril is projected onto $R_{cm}^{P_1P_2}$ and $Q_{tot}$ and is given by

$$F(R_{cm}^{P_1P_2}, Q_{tot}) = -k_B T \log P(R_{cm}^{P_1P_2}, Q_{tot}), \tag{S23}$$

where $P(R_{cm}^{P_1P_2}, Q_{tot})$ is the time independent joint probability distribution computed over the simulation trajectory using the equation

$$P(R_{cm}^{P_1P_2\prime}, Q'_{tot}) = \frac{\displaystyle\int_{t_{min}}^{t_{max}} d\tau \exp\left[\beta\big(V^b(R_{cm}^{P_1P_2}(\tau), Q_{tot}(\tau), \tau\big) - c(\tau)\big)\right] \delta(R_{cm}^{P_1P_2}(\tau) - R_{cm}^{P_1P_2\prime})\delta(Q_{tot}(\tau) - Q'_{tot})}{\displaystyle\int_{t_{min}}^{t_{max}} d\tau \exp\left[\beta\big(V^b(R_{cm}^{P_1P_2}(\tau), Q_{tot}(\tau), \tau) - c(\tau)\big)\right]}, \tag{S24}$$

where $t_{min}$ and $t_{max}$ are the lower and upper limits of the time scale where a local convergence is achieved along the collective variables $R_{cm}^{P_1P_2}$ and $Q_{tot}$, $c(t)$ is the time dependent bias offset, which estimates the local convergence in the CV space and is given by

$$c(t) = \frac{1}{\beta} \log \frac{\displaystyle\int dR_{cm}^{P_1P_2}\ dQ_{tot} \exp\left[-\beta F^{met}(R_{cm}^{P_1P_2}, Q_{tot})\right]}{\displaystyle\int dR_{cm}^{P_1P_2}\ dQ_{tot} \exp\left[-\beta\big(F^{met}(R_{cm}^{P_1P_2}, Q_{tot}) + V^b(R_{cm}^{P_1P_2}, Q_{tot}, t)\big)\right]}, \tag{S25}$$

where $F^{met}(R_{cm}^{P_1P_2}, Q_{tot})$ is computed using Tiwary-Parinello time independent free energy estimator,[18]

$$\beta F^{met}(R_{cm}^{P_1P_2}, Q_{tot}) = -\beta V^b(R_{cm}^{P_1P_2}, Q_{tot}, t) + \log \int dR_{cm}^{P_1P_2} dQ_{tot} \exp\left[\beta V^b(R_{cm}^{P_1P_2}, Q_{tot}, t)\right]. \tag{S26}$$

We constructed the free energy surface in the absence of the cosolvent, $[C] = 0$ M, using the Hamiltonian given by eq. S14. The effect of cosolvent on the free energy surface is computed by treating the cosolvent contribution term, $\Delta G_{tr}(\{\mathbf{r}\}, [C])$, in the Hamiltonian given by eq. S20 as a perturbation. The probability distribution $P(R_{cm}^{P_1P_2}, Q_{tot}, [C])$ in a cosolvent solution with concentration $[C]$ is given by[18]

$$P(R_{cm}^{P_1P_2\prime}, Q'_{tot}, [C]) = \frac{\displaystyle\int_{t_{min}}^{t_{max}} d\tau \exp\left[\beta\left(V^b(R_{cm}^{P_1P_2}(\tau), Q_{tot}(\tau), \tau) - c(\tau) - \Delta G_{tr}(\{\mathbf{r}\}, [C])\right)\right] \times \delta(R_{cm}^{P_1P_2}(\tau) - R_{cm}^{P_1P_2\prime})\delta(Q_{tot}(\tau) - Q'_{tot})}{\displaystyle\int_{t_{min}}^{t_{max}} d\tau \exp\left[\beta\left(V^b(R_{cm}^{P_1P_2}(\tau), Q_{tot}(\tau), \tau) - c(\tau) - \Delta G_{tr}(\{\mathbf{r}\}, [C])\right)\right]}, \tag{S27}$$

and the free energy surface $F(R_{cm}^{P_1P_2}, Q_{tot}, [C])$ in a cosolvent solution with concentration $[C]$ is given by

$$F\left(R_{cm}^{P_1P_2}, Q_{tot}, [C]\right) = -k_B T \log P\left(R_{cm}^{P_1P_2}, Q_{tot}, [C]\right). \tag{S28}$$

#### Free Energy Surface for the Domain Swapped Dimerization of cSrc-SH3

To compute the free energy of cSrc-SH3 domain swapped dimerization, we computed the time-independent probability distributions in each umbrella sampling window $h$ from the metadynamics simulations using the equations

$$P_h(Q'_{inter}, Q'_{intra,dim}, R'_{cm}) = \frac{\displaystyle\int_{t_{min}}^{t_{max}} d\tau \exp\left[\beta\left(V_h^b(Q_{inter}(\tau), Q_{intra,dim}(\tau), \tau) - c_h(\tau)\right)\right] \times \delta(Q_{inter}(\tau) - Q'_{inter})\delta(Q_{intra,dim}(\tau) - Q'_{intra,dim})\delta(R_{cm}(\tau) - R'_{cm})}{\displaystyle\int_{t_{min}}^{t_{max}} d\tau \exp\left[\beta\left(V_h^b(Q_{inter}(\tau), Q_{intra,dim}(\tau), \tau) - c(\tau)\right)\right]}, \tag{S29}$$

and

$$P_h(Q'_{inter}, Q'_{intra,dim}, Q'_{intra,mon}, R'_{cm}) = \frac{\displaystyle\int_{t_{min}}^{t_{max}} d\tau \exp\left[\beta\left(V_h^b(Q_{inter}(\tau), Q_{intra,dim}(\tau), \tau) - c_h(\tau)\right)\right] \times \delta(Q_{inter}(\tau) - Q'_{inter})\delta(Q_{intra,dim}(\tau) - Q'_{intra,dim})\delta(Q_{intra,mon}(\tau) - Q'_{intra,mon})\delta(R_{cm}(\tau) - R'_{cm})}{\displaystyle\int_{t_{min}}^{t_{max}} d\tau \exp\left[\beta\left(V_h^b(Q_{inter}(\tau), Q_{intra,dim}(\tau), \tau) - c(\tau)\right)\right]}. \tag{S30}$$

In the above equations, $c_h(t)$ is the time dependent bias offset in the umbrella sampling window $h$, which is computed using the equation[9,18]

$$c_h(t) = \frac{1}{\beta} \log \frac{\displaystyle\int dQ_{inter} dQ_{intra,dim} \exp\left[-\beta F_h^{met}(Q_{inter}, Q_{intra,dim})\right]}{\displaystyle\int dQ_{inter} dQ_{intra,dim} \exp\left[-\beta\left(F_h^{met}(Q_{inter}, Q_{intra,dim}) + V_h^b(Q_{inter}, Q_{intra,dim}, t)\right)\right]}, \tag{S31}$$

and $F_h^{met}(Q_{inter}, Q_{intra,dim})$ is the time independent free energy for cSrc-SH3 dimerization in the umbrella sampling window $h$, and it is computed using the Tiwary-Parrinello estimator[18] given by

$$\beta F_h^{met}(Q_{inter}, Q_{intra,dim}) = -\beta V_h^b(Q_{inter}, Q_{intra,dim}, t) + \log \int dQ_{inter} dQ_{intra,dim} \exp\left[\beta V_h^b(Q_{inter}, Q_{intra,dim}, t)\right]. \tag{S32}$$

To compute the unbiased probability distributions, $P(Q_{inter}, Q_{intra,dim}, R_{cm})$ and $P(Q_{inter}, Q_{intra,dim}, Q_{intra,mon}, R_{cm})$, we removed the umbrella bias $W_h^b(R_{cm})$ and combined probability distributions, $P_h(Q_{inter}, Q_{intra,dim}, R_{cm})$, and $P_h(Q_{inter}, Q_{intra,dim}, Q_{intra,mon}, R_{cm})$ from all the umbrella sampling windows using weighted histogram analysis method (WHAM)[19] equations given by

$$P(Q_{inter}, Q_{intra,dim}, R_{cm}) = \frac{\displaystyle\sum_{h=1}^{M} n_h P_h(Q_{inter}, Q_{intra,dim}, R_{cm})}{\displaystyle\sum_{h=1}^{M} n_h \exp[\beta f_h] \exp[-\beta W_h^b(R_{cm})]}. \tag{S33}$$

$$\exp\big[-\beta f_h\big] = \int dQ_{inter} dQ_{intra,dim} dR_{cm} \exp[-\beta W_h^b(R_{cm})] P(Q_{inter}, Q_{intra,dim}, R_{cm}) \tag{S34}$$

and

$$P(Q_{inter}, Q_{intra,dim}, Q_{intra,mon}, R_{cm}) = \frac{\displaystyle\sum_{h=1}^{M} n_h P_h(Q_{inter}, Q_{intra,dim}, Q_{intra,mon}, R_{cm})}{\displaystyle\sum_{h=1}^{M} n_h \exp[\beta f_h] \exp[-\beta W_h^b(R_{cm})]}, \tag{S35}$$

$$\exp\big[-\beta f_h\big] = \int \begin{matrix} dQ_{inter} dQ_{intra,dim} dQ_{intra,mon} dR_{cm} \exp[-\beta W_h^b(R_{cm})] \\ \times P(Q_{inter}, Q_{intra,dim}, Q_{intra,mon}, R_{cm}) \end{matrix}, \tag{S36}$$

where $n_h$ is the number of configurations visited in $h^{th}$ umbrella sampling window, and $M$ is the total number of umbrella windows.

The free energy surface for dimer formation projected onto the CVs, $Q_{inter}$ and $Q_{intra,dim}$ is computed using the equation

$$\begin{aligned} F(Q_{inter}, Q_{intra,dim}) &= -k_B T \log P(Q_{inter}, Q_{intra,dim}) \\ &= -k_B T \log \int dR_{cm} P(Q_{inter}, Q_{intra,dim}, R_{cm}), \end{aligned} \tag{S37}$$

and similarly we constructed $F(Q_{inter}, Q_{intra,mon})$ using the equation

$$\begin{aligned} F(Q_{inter}, Q_{intra,mon}) &= -k_B T \log P(Q_{inter}, Q_{intra,mon}) \\ &= -k_B T \log \int dQ_{intra,dim} dR_{cm} P(Q_{inter}, Q_{intra,mon}, Q_{intra,dim}, R_{cm}). \end{aligned} \tag{S38}$$

The effect of cosolvent with concentration $[C]$ on the probability distributions and the free energy surface for dimer formation in the umbrella sampling window $h$ is given by

$$P_h(Q'_{inter}, Q'_{intra,dim}, Q'_{intra,mon}, R'_{cm}, [C]) = \frac{\displaystyle\int_{t_{min}}^{t_{max}} d\tau \exp\left[\beta\big(V_h^b(Q_{inter}(\tau), Q_{intra,dim}(\tau)) - c_h(\tau) - \Delta G_{tr,h}(\{\mathbf{r}\}, [C])\big]\right. \times \delta(Q_{inter}(\tau) - Q'_{inter})\delta(Q_{intra,dim}(\tau) - Q'_{intra,dim})\delta(Q_{intra,mon}(\tau) - Q'_{intra,mon})\delta(R_{cm}(\tau) - R'_{cm})}{\displaystyle\int_{t_{min}}^{t_{max}} d\tau \exp\left[\beta\big(V_h^b(Q_{inter}(\tau), Q_{intra,dim}(\tau), \tau) - c(\tau) - \Delta G_{tr,h}(\{\mathbf{r}\}, [C])\big)\right]}, \tag{S39}$$

and

$$F_h(Q_{inter}, Q_{intra,mon}, [C]) = -k_BT \log\left(\int dQ_{intra,dim} dR_{cm} P_h\big(Q_{inter}, Q_{intra,dim}, Q_{intra,mon}, R_{cm}, [C]\big)\right). \tag{S40}$$

In the umbrella sampling window, where the monomers are restrained to large separations ($R_{cm}$ = 32 Å), the dimer formation is not possible and we only observe multiple folding-unfolding transitions between the cSrc-SH3 monomers, and the free energy for a monomer folding and unfolding transition is computed using the equation

$$F_h(Q_{intra,mon}, [C]) = -k_BT \log \int dQ_{inter} dQ_{intra,dim} dR_{cm} P\big(Q_{inter}, Q_{intra,dim}, Q_{intra,mon}, R_{cm}, [C]\big). \tag{S41}$$

### References


(1) Hyeon, C.; Dima, R. I.; Thirumalai, D. Pathways and kinetic barriers in mechanical unfolding and refolding of RNA and proteins. *Structure* **2006**, *14*, 1633–1645.

(2) Liu, Z.; Reddy, G.; O'Brien, E. P.; Thirumalai, D. Collapse kinetics and chevron plots from simulations of denaturant-dependent folding of globular proteins. *Proc. Natl. Acad. Sci. U. S. A.* **2011**, *108*, 7787–7792.

(3) Yu, H.; Rosen, M. K.; Schreiber, S. L. 1H and 15N assignments and secondary structure

of the Src SH3 domain. *FEBS Lett.* **1993**, *324*, 87–92.

(4) Camara-Artigas, A.; Martin-Garcia, J. M.; Morel, B.; Ruiz-Sanz, J.; Luque, I. Intertwined dimeric structure for the SH3 Domain of the c-Src tyrosine kinase induced by polyethylene glycol binding. *FEBS Lett.* **2009**, *583*, 749–753.

(5) Paravastua, A. K.; Leapman, R. D.; Yau, W. M.; Tycko, R. Molecular structural basis for polymorphism in alzheimer's beta-amyloid fibrils. *Proc. Natl. Acad. Sci. USA* **2008**, *105*, 18349–18354.

(6) Humphrey, W.; Dalke, A.; Schulten, K. VMD: Visual molecular dynamics. *J. Mol. Graph.* **1996**, *14*, 33–38.

(7) Betancourt, M.; Thirumalai, D. Pair potentials for protein folding: choice of reference states and sensitivity of predicted native states to variations in the interaction schemes. *Prot. Sci.* **1999**, *8*, 361–369.

(8) Laio, A.; Parrinello, M. Escaping free-energy minima. *Proc. Natl. Acad. Sci. U. S. A.* **2002**, *99*, 12562–12566.

(9) Awasthi, S.; Kapil, V.; Nair, N. N. Sampling free energy surfaces as slices by combining umbrella sampling and metadynamics. *J. Comput. Chem.* **2016**, *37*, 1413–1424.

(10) Veitshans, T.; Klimov, D.; Thirumalai, D. Protein folding kinetics: Timescales, pathways and energy landscapes in terms of sequence-dependent properties. *Fold Des* **1997**, *2*, 1–22.

(11) Swope, W.; Andersen, H.; Berens, P.; Wilson, K. A computer simulation method for the calculation of equilibrium constants for the formation of physical clusters of molecules: application to small clusters. *J. Chem. Phys.* **1982**, *76*, 637–649.

(12) O'Brien, E. P.; Ziv, G.; Haran, G.; Brooks, B. R.; Thirumalai, D. Effects of denaturants

and osmolytes on proteins are accurately predicted by the molecular transfer model. *Proc. Natl. Acad. Sci. U. S. A.* **2008**, *105*, 13403–13408.

(13) Liu, Z.; Reddy, G.; Thirumalai, D. Theory of the molecular transfer model for proteins with applications to the folding of the Src-SH3 domain. *J. Phys. Chem. B* **2012**, *116*, 6707–6716.

(14) Auton, M.; Bolen, D. W. Additive transfer free energies of the peptide backbone unit that are independent of the model compound and the choice of concentration scale. *Biochemistry* **2004**, *43*, 1329–1342.

(15) Reddy, G.; Liu, Z.; Thirumalai, D. Denaturant-dependent folding of GFP. *Proc. Natl. Acad. Sci. U. S. A.* **2012**, *109*, 17832–17838.

(16) Auton, M.; Bolen, D. W. Predicting the energetics of osmolyte-induced protein folding/unfolding. *Proc. Natl. Acad. Sci. U. S. A.* **2005**, *102*, 15065–15068.

(17) O'Brien, E. P.; Brooks, B. R.; Thirumalai, D. Molecular origin of constant m-Values, denatured state collapse, and residue-dependent transition midpoints in globular proteins. *Biochemistry* **2009**, *48*, 3743–3754.

(18) Tiwary, P.; Parrinello, M. A time-independent free energy estimator for metadynamics. *J. Phys. Chem. B* **2015**, *119*, 736–742.

(19) Kumar, S.; J.M., R.; Bouzida, D.; Swendsen, R.; Kollman, P. The weighted histogam analysis method for free-energy calculations on biomolecules. 1. The method. *J. Comput. Chem.* **1992**, *13*, 1011–1021.

Table S1: Energy function parameters for the SOP-SC model

| Parameters | cSrc-SH3 | $A\beta$ |
|---|---|---|
| $R_o$ | 2.0 | 2.0 |
| $k$ | 20 kcal/(mol. Å$^2$) | 20 kcal/(mol. Å$^2$) |
| $R_c$ | 8 Å | 8 Å |
| $r_0$ | 2 Å | 2 Å |
| $\epsilon_h^{bb}$ | 0.55 kcal/mol | 0.375 kcal/mol |
| $\epsilon_h^{bs}$ | 0.4 kcal/mol | 0.375 kcal/mol |
| $\epsilon_l$ | 1.0 kcal/mol | 1.0 kcal/mol |
| $\sigma^{bb}$ | 3.8 Å | 3.8 Å |
| $k_h$ | 2.0 kcal/(mol. Å$^2$) | 5.0 kcal/(mol. Å$^2$) |
| $k_{harm}$ | - | 15.0 kcal/(mol. Å$^2$) |

Table S2: SOP-SC model parameters for cSrc-SH3

| Parameters | cSrc-SH3 |
|---|---|
| $N_B$ | 111 |
| $N_N^{bb}$ | 151 |
| $N_N^{bs}$ | 350 |
| $N_N^{ss}$ | 144 |
| $N_N^{bb}(M_1, M_2)$ | 133 |
| $N_N^{bs}(M_1, M_2)$ | 295 |
| $N_N^{ss}(M_1, M_2)$ | 163 |
| $N_{NN}$ | 5241 |
| $N_{NN}(M_1, M_2)$ | 11953 |
| $N_{ang}^{bb}$ | 54 |
| $N_{ang}^{bs}$ | 110 |

Table S3: SOP-SC model parameters for $A\beta$

| Parameters | $A\beta$ |
|---|---|
| $N_B$ | 63 |
| $N_N^{bb}$ | 1 |
| $N_N^{bs}$ | 60 |
| $N_N^{ss}$ | 31 |
| $N_N^{bb}(P_1, P_2)$ | 100 |
| $N_N^{bs}(P_1, P_2)$ | 206 |
| $N_N^{ss}(P_1, P_2)$ | 90 |
| $N_{NN}$ | 1738 |
| $N_{NN}(P_1; P_2, P_3, P_4)$ | 20084 |
| $N_{ang}^{bb}$ | 30 |
| $N_{ang}^{bs}$ | 62 |

Table S4: Side-chain radii of amino acids

| Residue | Radius (Å) |
|---|---|
| Gly | 0.5 |
| Ala | 2.52 |
| Val | 2.93 |
| Leu | 3.09 |
| Ile | 3.09 |
| Met | 3.09 |
| Phe | 3.18 |
| Pro | 2.78 |
| Ser | 2.59 |
| Thr | 2.81 |
| Asn | 2.84 |
| Gln | 3.01 |
| Tyr | 3.23 |
| Trp | 3.39 |
| Asp | 2.79 |
| Glu | 2.96 |
| Hsd | 3.04 |
| Lys | 3.18 |
| Arg | 3.28 |
| Cys | 2.74 |

Table S5: Metadynamics parameters for $A\beta$ and cSrc-SH3 aggregation

| parameters | $A\beta$ | cSrc-SH3 |
|---|---|---|
| $T\ (K)$ | 350 | 353 |
| $\tau$ | 1000 | 1000 |
| $w\ (kcal/mol)$ | 0.05 | 0.05 |
| $\delta R_{cm}^{P_1P_2}$ (Å) | 5 | - |
| $\delta Q_{tot}$ | 10 | - |
| $\delta Q_{inter}$ | - | 10 |
| $\delta Q_{intra,dim}$ | - | 10 |

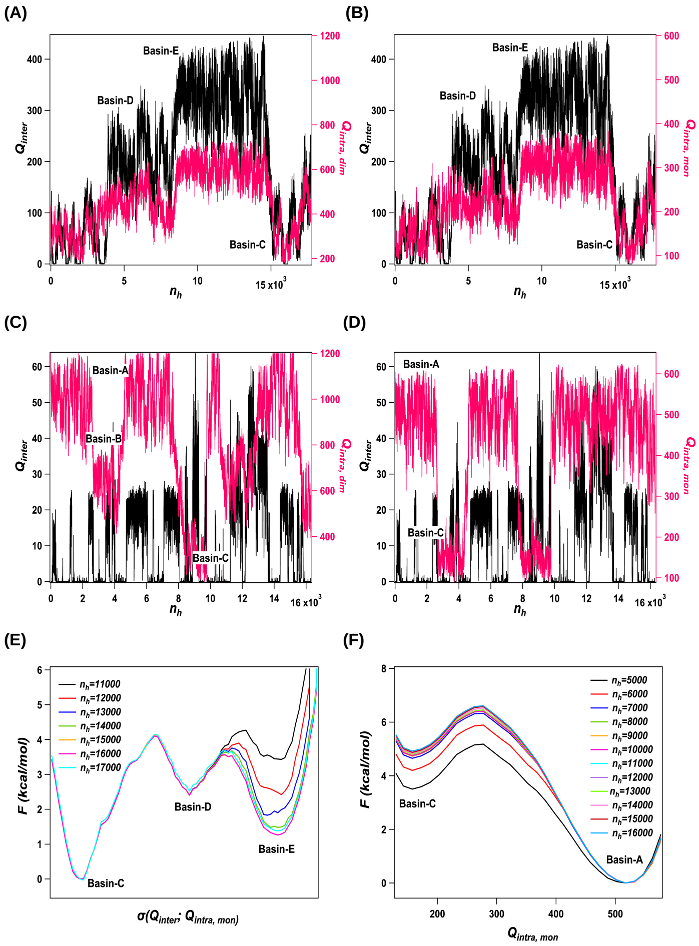


Figure S1: Convergence of the free energy surface for cSrc-SH3 domain swapped dimer formation computed using metadynamics simulations. (A) Sampling of the collective variables (CVs) $Q_{inter}$ and $Q_{intra,dim}$ as a function of the number of Gaussian hills ($n_h$) added during the metadynamics simulation in the umbrella sampling window where $R_{cm}$ is restrained at 4.5 Å. (B) Sampling of $Q_{inter}$ and $Q_{intra,mon}$ are plotted as a function of $n_h$ in the umbrella sampling window where $R_{cm}$ is restrained at 4.5 Å. In this umbrella sampling window, metadynamics simulation samples protein conformations in three basins corresponding to both the cSrc-SH3 monomers unfolded (basin-C), partially formed dimers (basin-D), and domain swapped dimers (basin-E). (C) Sampling of $Q_{inter}$ and $Q_{intra,dim}$ as a function of $n_h$ in the umbrella sampling window where $R_{cm}$ is restrained at 32 Å. (D) Sampling of $Q_{inter}$ and $Q_{intra,mon}$ as a function of $n_h$ in the umbrella sampling window where $R_{cm}$ is restrained at 32 Å. At this value of $R_{cm}$, the formation of domain swapped dimer is not feasible. The metadynamics simulations in this umbrella sampling window mainly sample protein conformations in three basins corresponding to both the monomers folded (basin-A), one monomer folded and the other monomer unfolded (basin-B), and both the monomers unfolded (basin-C). (E) Free energy for dimer formation, $F(Q_{inter}, Q_{intra,mon} \mid R_{cm} = 4.5$ Å), is plotted as a function of the minimum energy pathway (MEP), $\sigma(Q_{inter}; Q_{intra,mon})$, constructed for the umbrella sampling window with a restraint on $R_{cm} = 4.5$ Å, for different $n_h$. Convergence of the metadynamics simulation is achieved after the addition of $\approx 13000$ hills. (F) $F(Q_{intra,mon} \mid R_{cm} = 32$ Å) as a function of $Q_{intra,mon}$ for different $n_h$ shows that metadynamics simulation converges in this window after the addition of $\approx 7000$ hills.

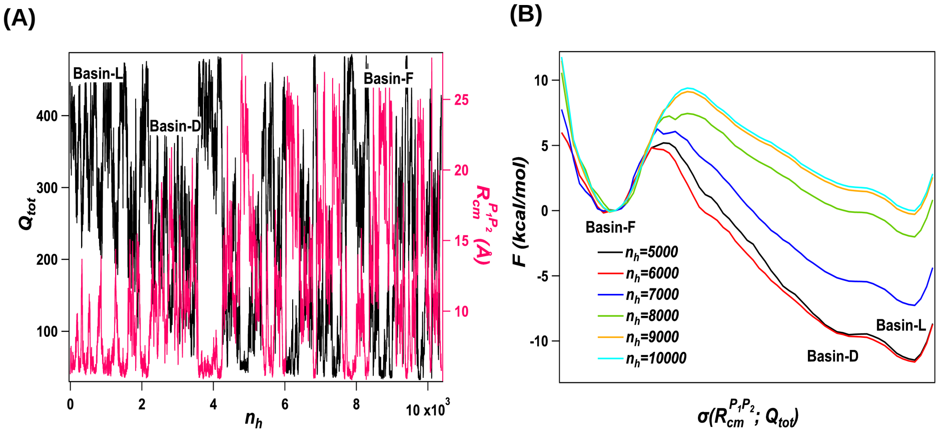


Figure S2: Convergence of the free energy surface associated with the dock-lock fibril growth step of the IDP $A\beta_{9-40}$. (A) Sampling of the CVs, $Q_{tot}$ and $R_{cm}^{P_1P_2}$, in the metadynamics simulations as a function of the number of Gaussian hills ($n_h$) added. Multiple transitions are observed among the three basins in the free energy surface, which correspond to the free state of the monomer in solution (basin F), the docked state where peptide in solution interacts with the fibril template (basin D), and the locked state where peptide becomes part of the fibril (basin L). (B) The free energy, $F(R_{cm}^{P_1P_2}, Q_{tot})$, projected onto the MEP, $\sigma(R_{cm}^{P_1P_2}; Q_{tot})$, shows that metadynamics simulations are converged after the addition of $\approx$ 8000 gaussians.

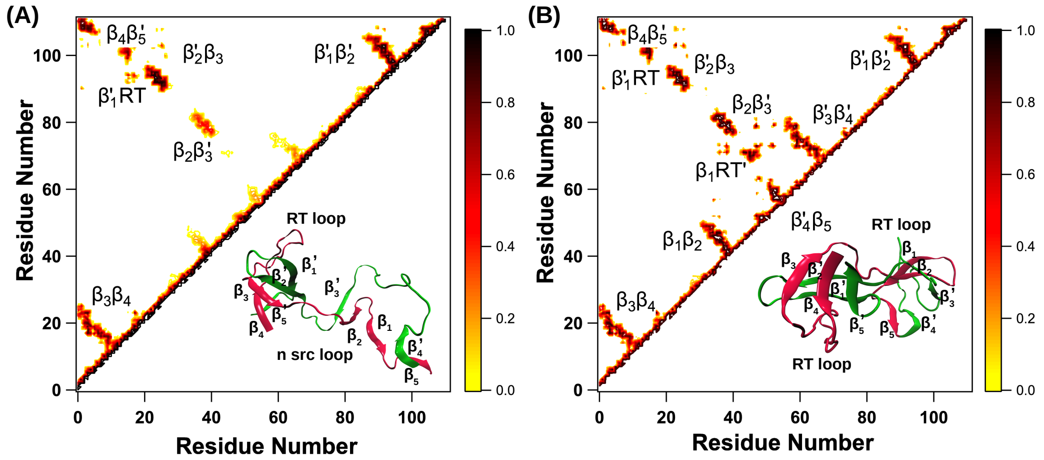


Figure S3: (A) Contact map of the protein conformations in the basin-D of cSrc-SH3 dimer formation. Nearly half of the contacts present in the domain swapped structure are formed in this intermediate state. (B) Contact map of the domain swapped structure of cSrc-SH3 obtained from the simulations.

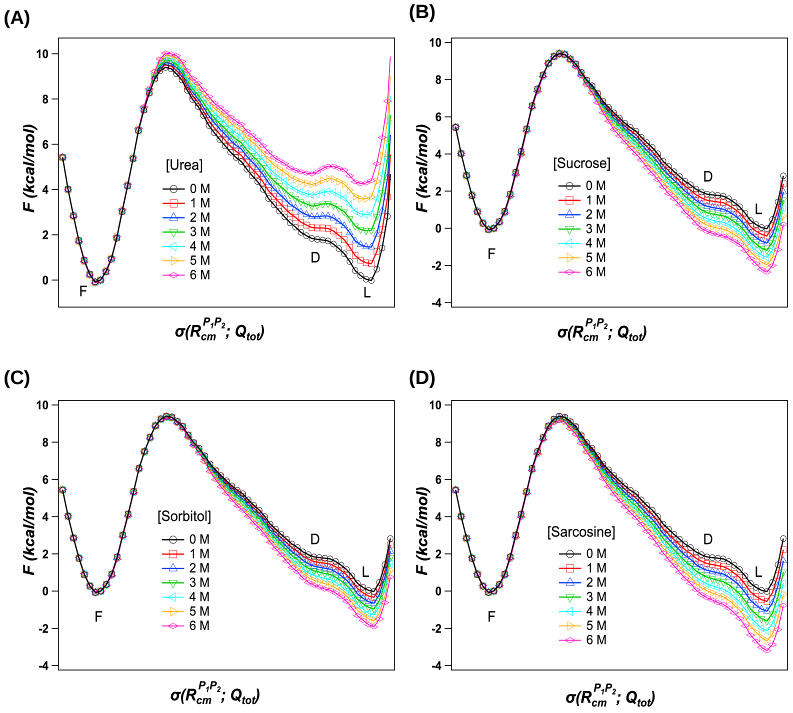


Figure S4: Effect of cosolvents on the free energy surface associated with the dock-lock fibril growth step of the IDP $A\beta_{9-40}$. Panels A, B, C and D show the effect of Urea, Sucrose, Sorbitol and Sarcosine, respectively on the free energy surface, $F(R_{cm}^{P_1P_2}, Q_{tot})$, projected onto the MEP, $\sigma(R_{cm}^{P_1P_2}; Q_{tot})$. Denaturants (Urea) stabilized the free monomeric state (basin F) while protective osmolytes (Sucrose, Sorbitol and Sucrose) stabilized the aggregated states (basins D and L). Denaturants also increased the energy barrier separating the free state from the docked and locked state.

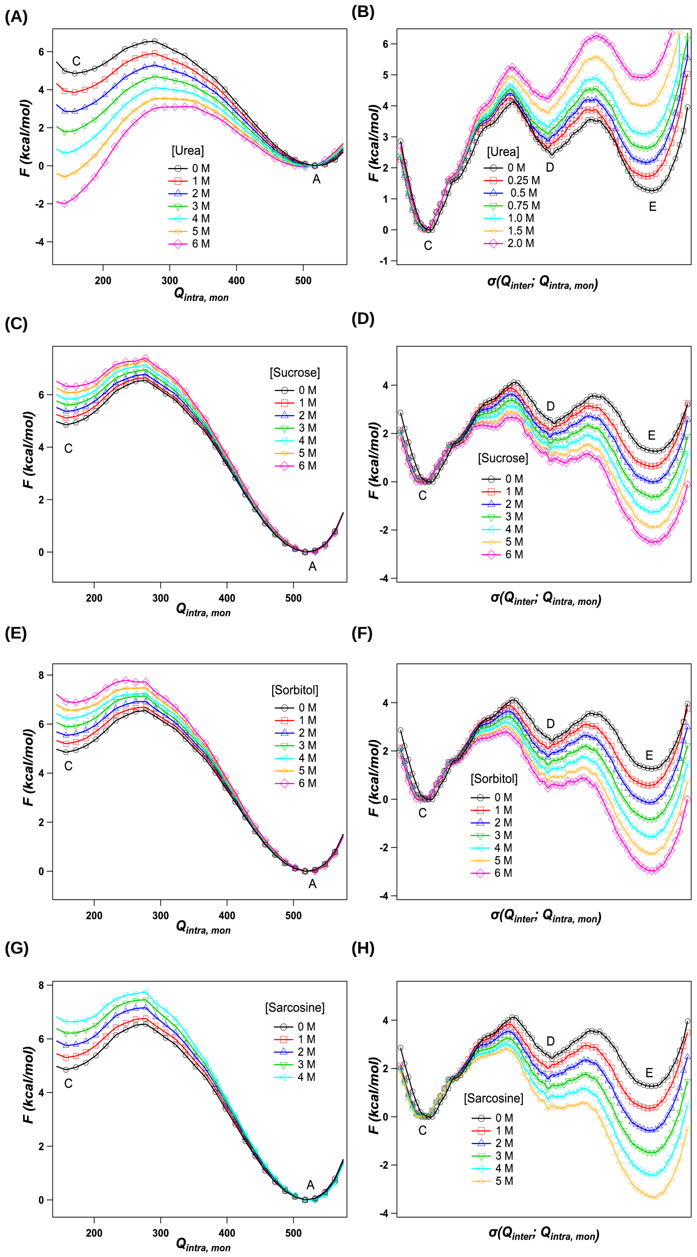


Figure S5: Effect of cosolvents on the free energy surface of cSrc-SH3 domain swapped dimer formation. Panels-A, C, E and G show the effect of Urea, Sucrose, Sorbitol and Sarcosine, respectively on the free energy surface, $F(Q_{intra,mon} \mid R_{cm} = 32\text{ Å})$ projected onto the CV, $Q_{intra,mon}$. When $R_{cm} = 32$ Å, domain swapped dimer formation is not possible, so we project the free energy only on $Q_{intra,mon}$. As expected, protective osmolytes stabilized the folded state (basin A), while denaturants stabilized the unfolded state (basin C). Denaturants decreased the energy barrier separating folded state from the unfolded states, while protective osmolytes increased the barrier. Panels-B, D, F and H show the effect of Urea, Sucrose, Sorbitol and Sarcosine, respectively on the free energy surface, $F(Q_{inter}, Q_{intra,mon} \mid R_{cm} = 4.5\text{ Å})$, projected onto the MEP, $\sigma(Q_{inter}; Q_{intra,mon})$. Denaturants increased the barrier height separating the unfolded state (basin - C) from the aggregated states (basins D and E), while protective osmolytes lowered the barrier height.
